# Erythropoietin restrains the inhibitory potential of interneurons in the mouse hippocampus

**DOI:** 10.1101/2023.09.29.560076

**Authors:** Yasmina Curto, Héctor Carceller, Patrycja Klimczak, Marta Perez-Rando, Qing Wang, Katharina Grewe, Riki Kawaguchi, Silvio Rizzoli, Daniel Geschwind, Klaus-Armin Nave, Vicent Teruel-Marti, Manvendra Singh, Hannelore Ehrenreich, Juan Nácher

## Abstract

Erythropoietin (EPO) aids in rectifying hippocampal transcriptional networks and synaptic structures of pyramidal lineages, thereby mitigating mood and cognition-associated disorders. An imminent conundrum is how EPO restores synapses by involving interneurons. By analyzing ∼ 12,000 single-nuclei transcriptomic data, we generated a comprehensive molecular atlas of hippocampal interneurons, resolved into 15 interneuron subtypes. Next, we studied molecular alterations upon recombinant human (rh)EPO and saw that gene expression changes relate to synaptic structure, trans-synaptic signaling and intracellular catabolic pathways. Putative ligand-receptor interactions between pyramidal and inhibitory neurons, regulating synaptogenesis, are altered upon rhEPO. An array of *in/ex vivo* experiments confirms that specific interneuronal populations exhibit reduced dendritic complexity, synaptic connectivity, and changes in plasticity-related molecules. Metabolism and inhibitory potential of interneuron subgroups are compromised, leading to greater excitability of pyramidal neurons. To conclude, improvement by rhEPO of neuropsychiatric phenotypes may partly owe to restrictive control over interneurons, facilitating re-connectivity and synapse development.

## INTRODUCTION

As the nomenclature reflects, the majority of studies on erythropoietin (EPO), a hypoxia-inducible growth factor, are dedicated to its hematopoietic function^1,2^. However, we and few other laboratories have long held the view that EPO also exerts potent, hematopoiesis-independent, neuroprotective, neuroregenerative and procognitive functions in the brain^3–9^. Using a medley of genomics/transcriptomics, functional assays, histology and behavioral readouts in rodents as experimental models, seminal studies have demonstrated beneficial effects of EPO and its receptor (EPOR) in postnatal hippocampal neurogenesis, ultimately resulting in improved cognitive performance, and thus stimulating multiple translational clinical trials^10–17^. Indeed, transgenic strategies demonstrated that constitutive overexpression of EPOR in principal neurons enhances cognitive functions in mice^18^. This improvement builds on the EPO-mediated maneuvering of neuronal differentiation trajectories, overpopulation of distinct pyramidal lineages, enhanced dendritic spine density and improved motor-cognitive execution, which is overlaid with induction of related gene regulatory networks^17,19–21^.

While the plasticity of excitatory hippocampal circuits bears the most of rhEPO-induced changes in cognition, pyramidal neurons from the CA1 region are accurately timed and synchronized by a rich diversity of GABAergic interneurons^22–26^. It is consensus that the different classes of interneurons interact with specific cellular domains of pyramidal neurons to establish the excitatory:inhibitory (E:I) balance, properly modulating cell firing, plasticity, network activity, metabolism, and trans-synapses. Like excitatory neurons, interneurons in the adult brain are capable of remodeling their structure and connectivity through changes in the expression of plasticity-related molecules, e.g. the polysialylated form of the neural cell adhesion molecule (polySia-NCAM), and specialized regions of the extracellular matrix denominated perineuronal nets (PNNs)^27–32^. How or to which extent interneurons remodel in response to rhEPO has remained enigmatic. Given the seminal papers in this field, it is imperative to determine the impact of rhEPO on interneurons to better resolve the mechanistic basis of cognitive advance. We wondered whether interneurons, subjected to rhEPO, undergo structural or physiological alterations, and molecular changes in gene regulatory networks, controlling crosstalk of pre- and post-synapses.

In the hippocampus, basket cells represent one of the main interneuronal subpopulations, given their abundance and strong inhibitory input on pyramidal neurons^22^. They extend their axonal arbor in the *stratum pyramidale* and vastly target the somata and proximal dendrites of pyramidal neurons^24,25,33^. This group is represented by the parvalbumin-positive (PVALB), fast-spiking and the regular-firing cholecystokinin-expressing (CCK) basket cells. The second largest group of hippocampal interneurons is formed by the dendrite-targeting interneurons, being the somatostatin (SST)-expressing O-LM (oriens/lacunosum-moleculare) cells, the main subpopulation in the CA1 region. They strongly inhibit the distal dendritic tuft of pyramidal neurons, usually in a feedback manner. The axons of SST O-LM cells arborize in the *stratum lacunosum-moleculare*, where they form a plexus of *en passant boutons* (*EPB*), which stablish synapses on dendrites of CA1 pyramidal cells^34,35^. Interestingly, SST O-LM interneurons exhibit dendritic spines that remain highly plastic during adulthood^36,37^.

Although mouse studies have carefully detailed the characteristics and events that precede the formation and survival of interneurons, molecular criteria to define interneuron subgroups within major lineages seem sparse^25,38–40^. This might either be due to their highly diverse and heterogeneous nature, or the conjecture that their specification is influenced by intrinsic or extrinsic factors^41–43^. High-resolution reference transcriptome of all hippocampal interneuron lineages following rhEPO treatment would greatly benefit the present study, but is currently lacking.

Even though the structural and molecular basis of interneurons in mammals is functionally similar, our understanding of their diversity and mechanisms of regulation in hippocampus remains limited^39,44^. Here, we first aimed to resolve the transcriptomic landscapes of interneuronal lineages in the murine hippocampus and then investigated alterations upon rhEPO to potentially reveal mechanistic details on its action. Identifying the key interneuron lineages and their modified cell-to-cell communication with pyramidal neurons should help to discern how synapses are renovated by rhEPO. With a strategic analysis of our single nuclei (sn)RNA-seq datasets^21^, we resolve 15 transcriptionally distinct interneuron lineages. Upon whittling the genes from the transcriptome of each lineage that are differentially expressed upon rhEPO, we show that EPO modulates the expression of genes involved in neurite morphology, synapses and metabolism in a lineage-specific manner. Also, we uncover the in-silico anatomical organization of pyramidal-interneuron connections mediated by ligand-receptor interactions that are re-bridged upon rhEPO. We overlaid our transcriptomics approach with robust *in/ex vivo* experiments to understand the impact of rhEPO over metabolism and plasticity of interneurons.

## RESULTS

### The snRNA-seq data analysis of hippocampal interneurons resolves 15 distinct lineages

A growing list of seminal studies has illustrated the enormous diversity of hippocampal interneurons. Accountability of rhEPO in altering the interneuronal transcriptome is still obscure^39,45^. Thus, we first classified all existing sub-lineages at single interneuron level. From the snRNA-seq of 12 hippocampi [6 each for rhEPO and placebo (PL) treatment], we obtained ∼ 108,000 single-nuclei transcriptomes grouped into 11 major cell types^21^. Among them, ∼16,000 cells (15%) nuclei were defined as interneurons, according to our cell-type classification and supported by their corresponding bonafide markers, allowing to identify interneuron subtypes.

Performing an unsupervised clustering using the most variable genes and graph-based learning on our snRNA-seq dataset^21^ (see methods), we identified numerous distinct clusters of interneurons (**Supplementary Figure 1A**). Clusters were obtained after a rigorous batch-control algorithm, preventing sample biases (**Supplementary Figure 1A**), and classified based on differential expression of transcriptomes. To ensure clusters to be homogeneously interneurons, expression of genes marking both interneuron and non-interneuron lineages were screened. Though the majority of clusters were interneurons, i.e. co-expressing *Gad1* and *Gad2* genes but not markers of other lineages, a few clusters expressed markers of microglia, oligodendrocytes or Cajal-Retzius cells (**Supplementary Figure 1B**). Although it is tempting to resolve ancestral and derived cells of these heterogeneous clusters, we removed them in a stepwise process (see methods) and focused on mature homogeneous interneurons and not the transitory or yet-to-be-committed cells (**Figure 1A**). These lineages expressed *Gad1* and *Gad2* at a higher level without displaying marker gene expression of other hippocampal lineages. For instance, *Tgfbr1* for microglia, *Gfap* for astrocytes, *Plp1* and *Pdgfra* for oligodendrocytes, *Bsg* for pericytes, *Flt1* for endothelium and *Slc17a6/Neurod6* for pyramidal neurons were tested in our classified interneuronal clusters. Again performing unsupervised clustering of the remaining ∼9000 nuclei^21^, we identified 15 distinct clusters (**Figure 1A-B**). These nuclei did not cluster by batch or samples, indicating a robust control of such effects (**Supplementary Figure 1A**). Instead, we classified these 15 clusters based on the combination of known, bonafide and discovered gene expression markers^39^ (**Figure 1C, Table S1**). Our data resolved the distinct transcriptome signature separating three types of SST, two CCK, two PVALB, two IS, two Ivy cells, and two distinct lineages termed Trilaminar and Cholinergic interneurons. We also observed two previously uncharacterized interneuronal populations that we classified based on top marker genes as *Nrg1/Ptprd* and *Zbtb20/Mgat4c* interneurons (**Figure 1A-C**). Collectively, we provide a transcriptome reference frame as valuable resource to interpret and analyze the diversity and heterogeneity of hippocampal interneurons via comprehensive survey of snRNA-seq data postdating rhEPO and PL treatment.

**Figure 1.**
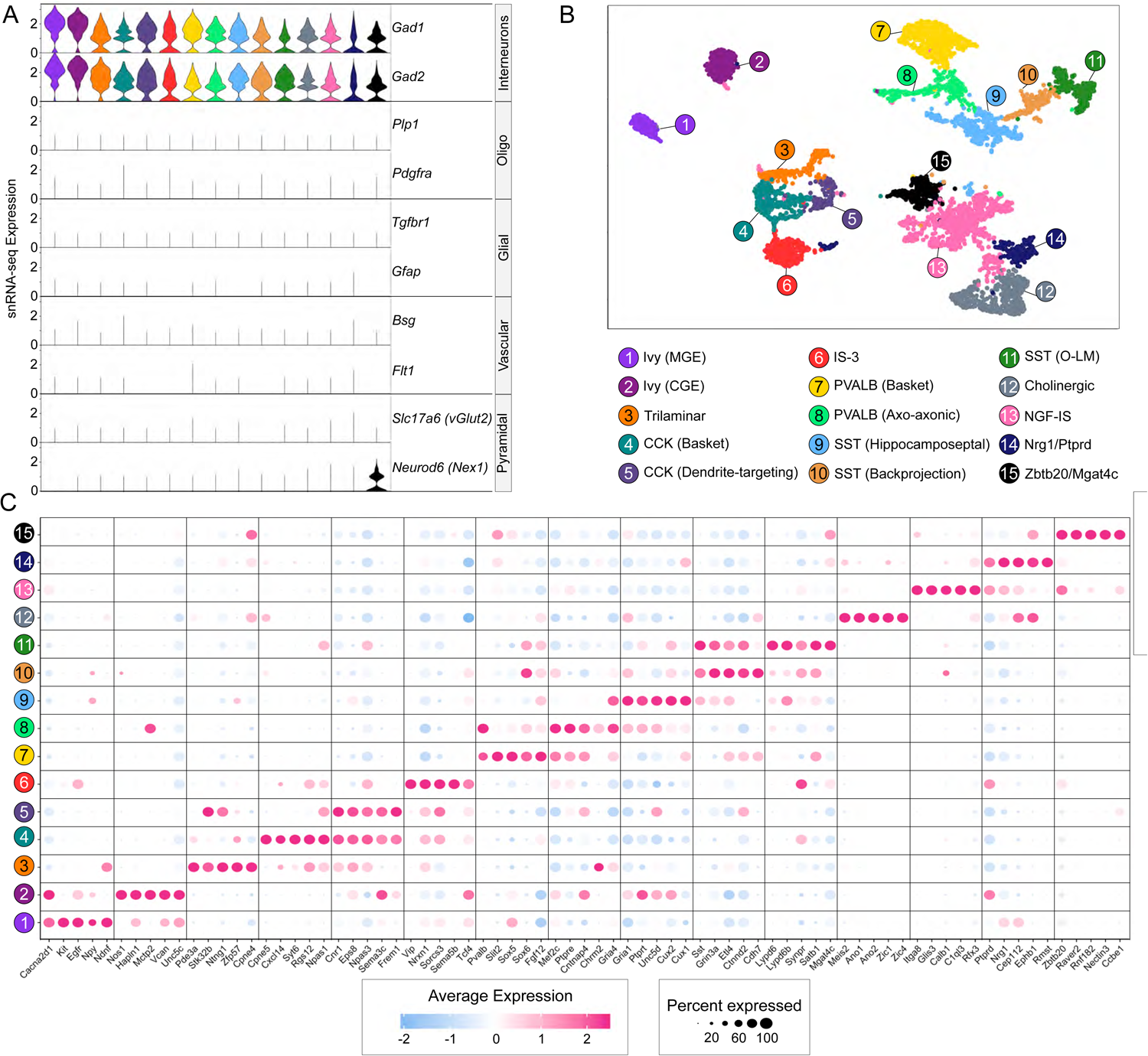
Single nuclei landscape of interneuronal lineages from 12 murine hippocampi. **A:** Violin plots illustrating the expression dynamics of cell type specific marker genes (two each) of interneurons, oligodendrocytes, glial (microglia and astrocytes), vascular (endothelial and pericytes) and pyramidal lineages across all obtained clusters. The 15 clusters are sublineages of mature interneurons, each cluster is color-coded as in Figure 1B. **B:** Two-dimensional Uniform Manifold Approximation Plot (UMAP) resolving ∼8,000 single nuclei into 15 distinct clusters, merged from 12 adult hippocampi of mice treated with either rhEPO(N=6) or PL(N=6). The clusters affiliate with interneuron subtypes, validated by testing co-expression of *Gad1* and *Gad2* genes (Figure 1A). Absence of transcriptional markers from the rest of hippocampal lineages was also confirmed. Colors indicate an unbiased classification of nuclei via graph-based clustering, where each dot represents a single nucleus. **C:** Dotplot illustrating cluster-specific expression of top marker genes for interneuronal populations identified from the above data. Cluster numbers derived from Figure 1B are given on the left of the plot. Marker gene names are presented at the bottom of the plot. Colors represent an average Log2 expression level scaled to the number of unique molecular identification (UMI) values in single nuclei. The color scale ranges from light blue to pink, corresponding to lower and to higher expression. Dot size is proportional to the percent of cells expressing the corresponding gene.

### rhEPO modulates gene expression of interneuronal subpopulations involved in E:I balance

The broader understanding of rhEPO mode of action warrants re-investigating the above identified interneuron lineages for differential expression of genes (DEGs) upon rhEPO. We first determined the aggregated expression of each gene from all nuclei within a lineage from rhEPO against PL samples, and then calculated the level of their differential expression using the recommended statistical set-up (see methods). Given their relevance in the mouse hippocampus, we curbed our DEGs analysis to SST (O-LM and backprojection), PVALB (basket and axo-axonic cells), and CCK (basket and dendrite-targeting) lineages. Using the statistical threshold of adjusted p-value <0.05, our analysis proclaims 1073 DEGs, accounting for all comparisons in these six lineages. While a few DEGs were shared between multiple lineages, most were unique to a particular lineage, suggesting that the impact of rhEPO on gene expression occurs in cell-type-specific manner (**Figure 2A, Table S2**).

**Figure 2.**
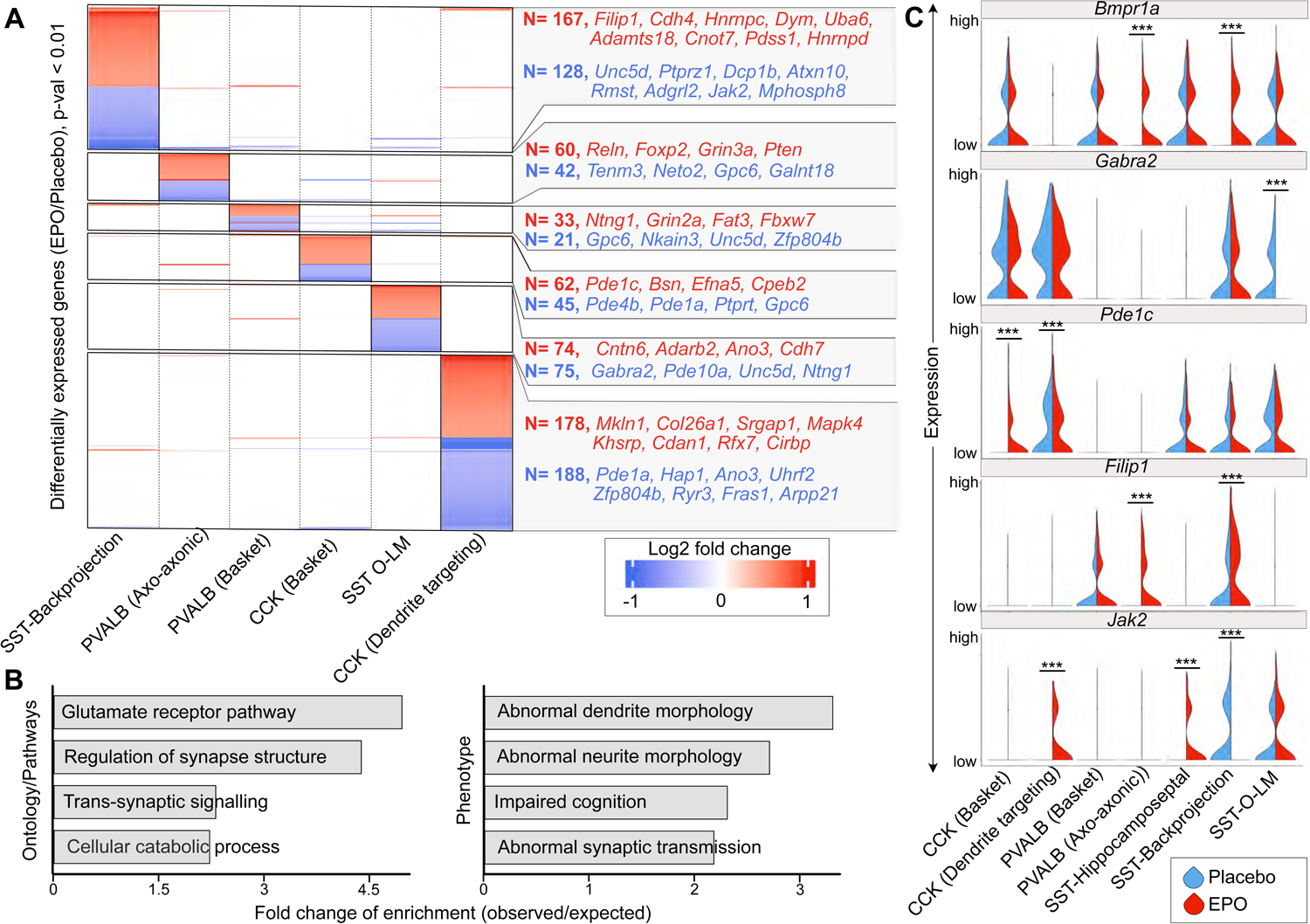
rhEPO modulates differential expression of synapse-associated genes in interneuron lineages. **A:** Heatmap representing the differential expression of genes between rhEPO and PL samples in individual lineages shown on the bottom. Only those genes with adjusted p-value <0.01 in any of the comparisons were used for plotting. The number of detected differentially expressed genes in each lineage is shown at the right side of the heatmap. The genes with insignificant differential expression are colorless in this figure. Annotation of gene names are the top DEGs in the given lineage (red, upregulated; blue, downregulated). **B:** Barplot showing the gene ontologies in which genes that are differentially expressed between rhEPO and PL cells are enriched. The strength of enrichment is labelled on the X-axis. **C:** Violin plots showing the expression distribution of *Bmpr1a, Gabra2, Pde1c, Filip1, and Jak2* genes between rhEPO and PL samples in a pairwise fashion among the individual lineages of PVALB, CCK and SST interneurons. Adjusted p-values are obtained by Benjamini-Hochberg (BH) correction.

To get a functional clue of the impact of rhEPO treatment, we conducted pathway and gene ontology analyses on the 1073 DEGs, applying KEGG databases and biological processes databases from Gene Ontology Consortium. These DEGs overrepresent genes related to glutamate receptors, synaptic structure, trans-synaptic signaling and intracellular catabolic pathways (**Figure 2B**). We further asked if these DEGs also overrepresent gene sets associated with documented phenotypes at molecular and individual levels upon rhEPO^10–17^. Expectedly, we found phenotypic association with dendrite morphology, synaptic transmission and cognition (**Figure 2B**), which are affirmed in previous studies to be positively regulated by rhEPO^12,17,19^.

Among the genes whose expression was toned down in rhEPO samples, we identified *Gabra2* and *Unc5d* in the SST O-LM lineage, whose repression might lead to loss of the inhibitory effect (**Figure 2A, and 2C**). On one hand, *Gabra2* keeps the inhibition of neurons intact at the synapse^46^; on the other hand, *Unc5d* facilitates migration of interneurons to the synapse^47^. It is compelling to hypothesize that repression of these two genes could be one of the many causes of higher excitability of pyramidal neurons in rhEPO subjects, as shown previously^21^. Among the upregulated genes in rhEPO samples, we noticed *Filip1* in both PVALB and SST lineages (**Figure 2C**). *Filip1* encodes a structural protein which shapes dendrite or neurite morphology and mediates neuronal migration^48,49^. Moreover, *Bmpr1a*, an essential receptor regulating downstream BMP signaling, is among the upregulated genes in both PVALB and SST lineages upon rhEPO (**Figure 2C**). Of note, recent genetic experiments have illustrated that *Bmpr1a* guides specific synaptic connectivity in PVALB interneurons^50,51^. In addition, we found few other genes with opposite trend of differential expression between sister lineages, also associated with synaptic pruning and maturation of interneurons. For instance, *Jak2*, known to prune inactive synapses^52,53^ appeared to be induced in SST-hippocamposeptal and CCK dendrite-targeting cells; whereas its expression disappeared in the SST-backprojection lineage upon rhEPO (**Figure 2C**). Similarly, *Pde1c*, a phosphodiesterase enzyme, mediating neuronal oxidative metabolism^54^, has gain and loss of expression in CCK basket and dendrite-targeting lineages, respectively, upon rhEPO (**Figure 2C**). Notably, on the list of DEGs, we also identifed multiple phosphodiesterase genes possessing oxidative metabolic properties, further corroborating our hypothesis that rhEPO can palliate metabolic dysregulations (**Figure 2A**). The above results point towards an interplay of rhEPO with interneuronal physiology, much more diverse than previously appreciated. It is enticing to hypothesize that the rhEPO-governed gene expression changes attenuate the interaction of interneurons with their target cells. This might underpin molecular criteria defining rhEPO-enforced interneuron plasticity, metabolism, and E:I balance.

### rhEPO modifies the cell-to-cell interactions between interneuronal and pyramidal lineages

So far, we demonstrated that the impact of rhEPO might be associated with mitigation of intracellular signaling cascades in interneurons. We hypothesized that this impact might often be due to a differential receptor-ligand activity. To systematically study the interactions between interneuronal and pyramidal lineages, we employed *LIANA* framework, a specialized tool for ligand-receptor analysis, using snRNA-seq data to infer cell-cell communication^55^. *LIANA* leverages the expression levels of genes overlaying with the CellPhoneDB repository of ligand–receptor pairs^56^, reckoning their architecture at subunit levels, including the secreted and cell-surface molecules. We first apprehended the diverse interneuronal and pyramidal lineages in a single data frame^21^. Then we asked what plausible interactions are altered significantly under the influence of rhEPO. Our cross-lineage comparison illustrates an array of interactions between interneurons and pyramidal neurons linegaes (**Supplementary Figure 2**, **Table S3**). It revealed various known and new ligand–receptor interactions. For instance, communication of mature CA1 pyramidal neurons with PVALB, CCK, and SST interneurons occurs in both rhEPO and PL samples through interactions of the secreted SLIT protein families binding to ROBO receptor families^57^ or between *Cadm1* (PVALB, CCK, and SST interneurons) and *Nectin3* (pyramidal mature neurons)^58^ (**Figure 3A-F**, **Supplementary Figure 2**, **Table S3**). Interactions between neuroligins (*Nlgn1/2/3*) and neurexins (*Nrxn1/2/3*), popular synaptic cell-adhesion molecules to establish the synapse^59^, were observed in all of the cross-lineage comparisons, supporting the fidelity of our data analysis. While we note that *Nrg3* is known to bind *Erbb3/4*^60^, which regulates E:I balance between pyramidal and PVALB interneurons^61^, our data show a highly significant binding between *Nrg3* and *Egfr* in PVALB axo-axonic lineage specifically (**Figure 3C**, **Supplementary Figure 2**). Because *Egfr* possesses a similar receptor structure as *Erbb3/4*^62^, it is plausible that *Nrg3* might serve *Egfr* as a ligand, but this mystery needs to be resolved in upcoming studies.

**Figure 3.**
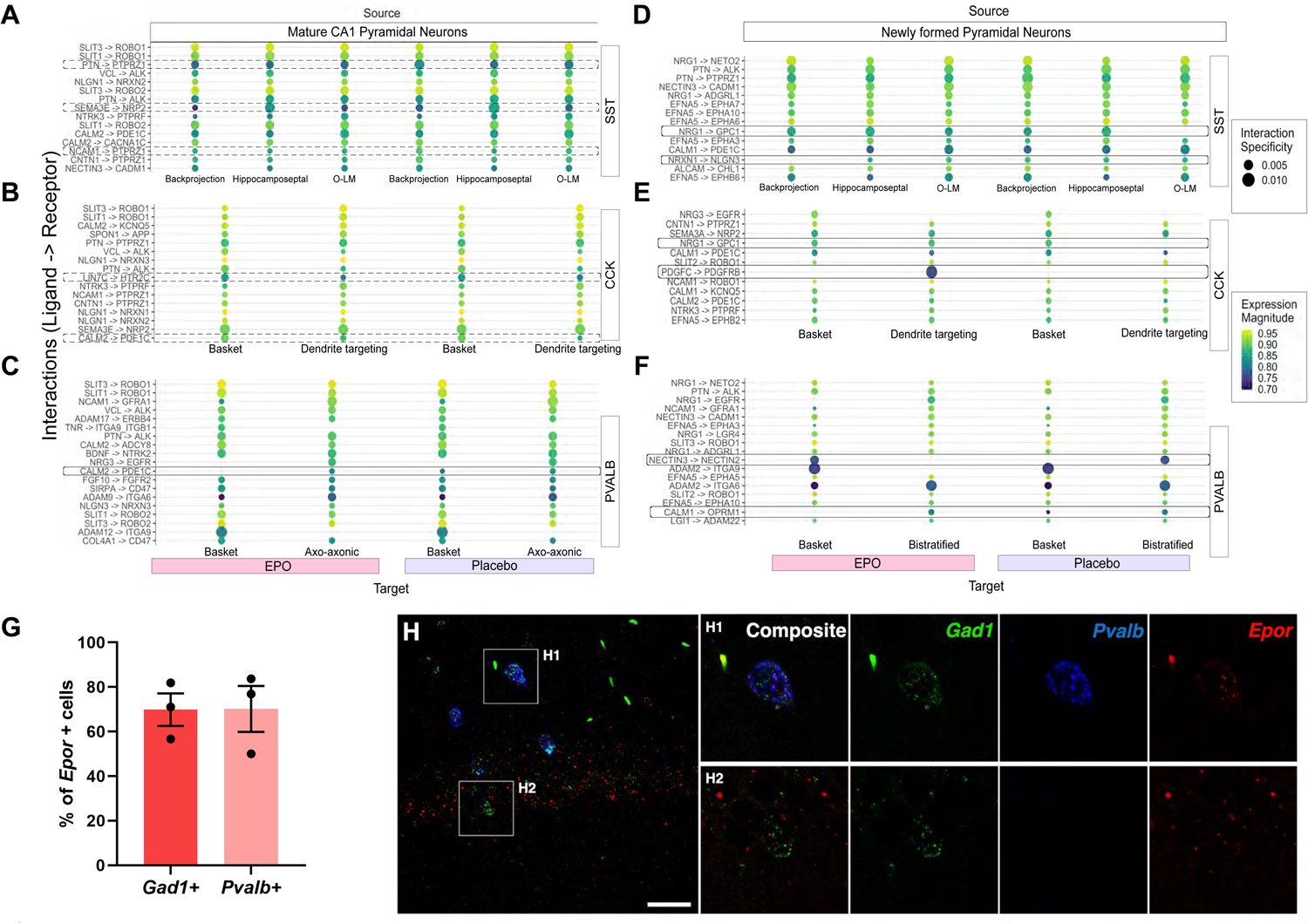
Modification by rhEPO of cell-to-cell interactions between interneuronal and pyramidal lineages and expression of EPOR in interneuronal subpopulations. Cell-to-cell communication analysis predicts source/target of cross-talks between pyramidal and interneuronal lineages among rhEPO and PL samples. These communications are demonstrated as an array of dotplots. The color scale ranges from dark blue to yellow-green, corresponding to lower and to higher co-expression magnitude. Dot size is proportional to the intensity of their interactions, specificity given in CellPhoneDB^56^. **A:** Baiting mature CA1 pyramidal lineages as a source, the dotplot shows the differential interaction of ligands *Ptn*, *Ncam1*, *Sema3e*, with *Ptprz1*, or *Nrp2* receptors, respectively, encoded by distinct lineages of SST interneurons. **B-C:** Similarly, while the *Calm2* interaction with *Pde1c* is diminished in CCK-dendrite targeting lineage, PVALB basket cells lack this interaction completely in rhEPO samples. **D:** The dotplot as above shows the differential interaction of newly formed pyramidal neurons with SST interneurons between rhEPO and PL samples. While *Nrg1-Gpc1* appears in O-LM, the *Nrxn1-Nlgn3* interaction is lost in the Backprojection lineages in rhEPO samples. E: *Pdgfc*-*Pdgfrb,* and *Nrg1-Gpc1* interactions emerge between CCK-dendrite targeting cells and newly formed pyramidal neurons in rhEPO samples. **F:** Curiously though, *Nectin3*-*Nectin2* interaction is shifted from PVALB bistratified cells to basket cells upon comparison of gain and loss of interactions between PL to rhEPO samples. However, the *Calm1*-*Oprm1* interaction is lost in basket cells of rhEPO samples. **G:** Graph showing the percentage of *Gad1*+ and *Pvalb*+ interneurons expressing *Epor* in CA1 hippocampal region. **H:** Representative FISH image from CA1 demonstrating *Epor* mRNA expression in *Gad1+* and *Pvalb+* interneurons. Magnification of focal planes show co-expression of *Epor* (red) with *Gad1* (green) and *Pvalb* (blue) (H1-H2). Scale bar: 35µm for overview in H and 16µm for focal planes.

We next assessed rhEPO-specific differences in the ligand-receptor profiles. While most of the interactions were similar in our catalogue of ligand-receptor repertoire, upon closer inspection, we observed a subtle loss and a gain of interaction in rhEPO samples. To elaborate, the *Calm2-Pde1c* pair, a known interaction partner^63^ was lost between PVALB basket cells and mature CA1 pyramidal neurons upon rhEPO, suggesting the attenuation of calcium-dependent signaling between these two lineages (**Figure 3C**). The loss of this interaction is consistent with the downregulation of a bunch of genes encoding phosphodiesterases in interneuron lineages. Furthermore, *Pdgfc-Pdgfrb* paired between newly formed pyramidal neurons and CCK-dendrite targeting interneurons was surfaced in rhEPO samples in contrary to PL (**Figure 3E**). Interestingly, this ligand-receptor is known to regulate neurogenesis, cell survival and synaptogenesis in hippocampus^64–66^, which supports the notion that EPO mediates hippocampal neurogenesis and synaptogenesis^17,19,21^. Together, this data supports a potential control of rhEPO over trans-synaptic signaling and E:I balance through tinkering specific cell-to-cell communications between interneurons and pyramidal lineages.

While the segregation of rhEPO and PL samples through DEGs and ligand-receptor communications in interneurons is compelling, it is merely based on expression at RNA level and there are certain limitations to the snRNA-seq data, as noted in multiple reports before. Despite these limitations, the above observations suggest that the alterations in interneuron physiology are acquired due to rhEPO treatment. To concur this, we inquired whether interneuronal populations express EPOR, which should establish that they can directly respond to rhEPO. We employed a highly sensitive fluorescence *in-situ* hybridization (FISH) method, able to detect *Epor* mRNA (**Figure 3G-H**). We analyzed the expression of *Epor* in GAD67+ (encoded by *Gad1*) cells, a typical GABAergic interneuronal marker, and PVALB+ interneurons (*Pvalb*), one of the key subpopulations of hippocampal interneurons. The analysis revealed 69.86±7.30% and 70.19±10.28% of GAD67*+* and PVALB*+* interneurons, respectively, co-expressing *Epor* in the hippocampal CA1 region (**Figure 3G**).

So far, the differential expression profiles and *Epor* expression seem to agree with the hypothesis that rhEPO modulates the interneuronal population; however, these findings still lack validation at cellular and physiological levels. We therefore performed a series of *in/ex vivo* experiments and specifically asked whether E:I balance, structural and synaptic properties, and metabolism of specific interneuronal subpopulations are changed upon rhEPO.

### rhEPO treatment decreases the structural complexity of specific hippocampal interneurons

Following the above leads, we analyzed - after 3-week PL/rhEPO treatment - structural changes in two major subpopulations of interneurons (**Figure 4A-B**): (i) GAD-EGFP positive interneurons, which mainly correspond to SST O-LM cells^37^ and (ii) PVALB interneurons (PV+).

**Figure 4.**
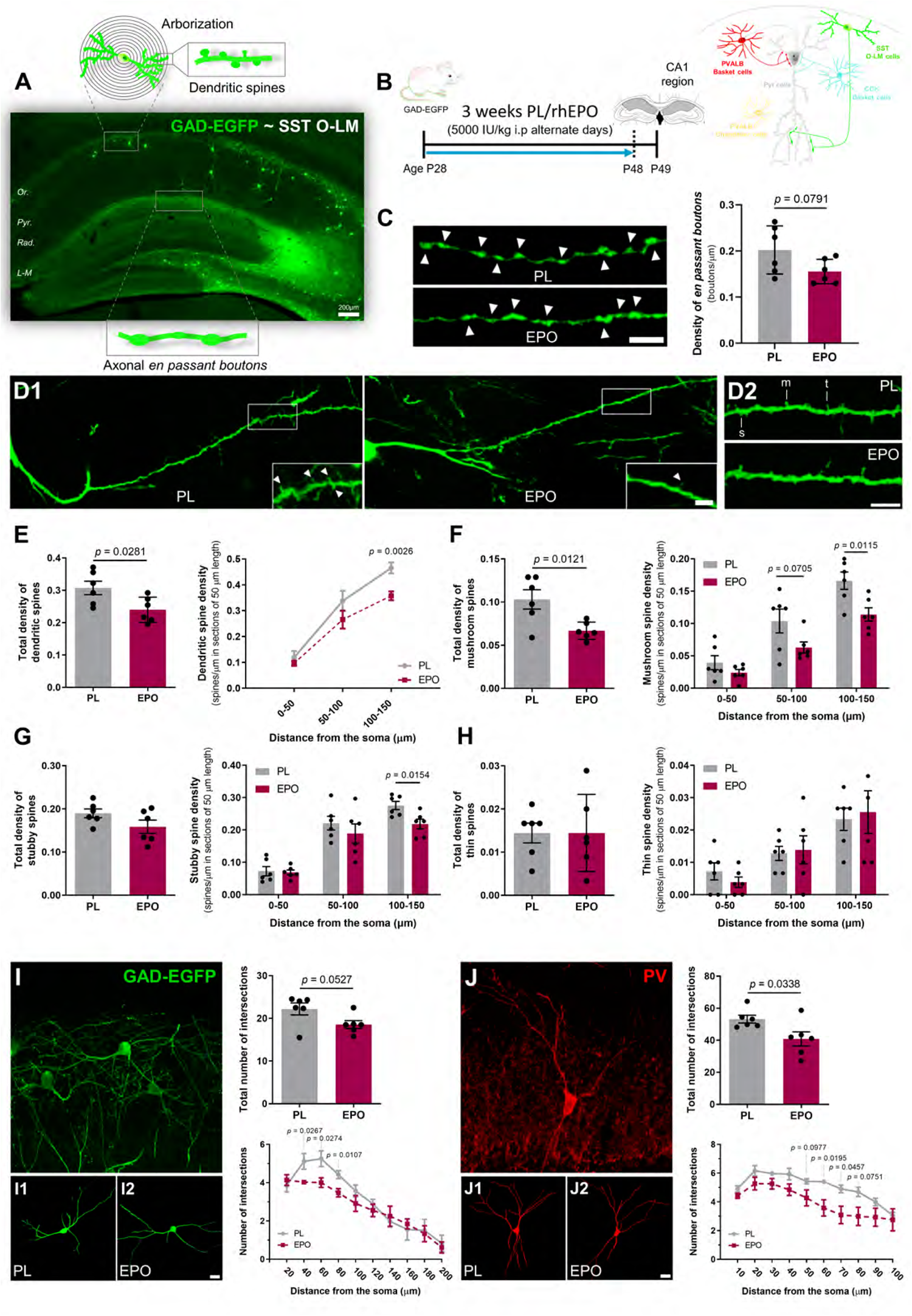
Treatment with rhEPO decreases the structural complexity of SST O-LM cells and PV-expressing interneurons in the hippocampal CA1 region. **A:** Representative hippocampal image of GAD-EGFP mice showing a schematic of different structural analyses performed. The majority of GAD-EGFP cells correspond to SST O-LM interneurons. **B:** Treatment scheme and hippocampal area of analysis (CA1 region) illustrated with different interneuronal subtypes and their communication with pyramidal neurons. **C:** Analysis of GAD-EGFP positive *EPB* (arrowheads) in the *stratum lacunosum-moleculare.* Confocal 2D projections show EGFP-positive axons of rhEPO and PL treated mice. Graph represents the changes in the density of *EPB,* expressed as boutons per micron. **D:** Analysis of dendritic spine density in GAD-EGFP expressing cells located in *stratum oriens.* Representative fluorescent images of spiny dendrites are shown for rhEPO and PL treated mice (D1). Dendritic spines are indicated by arrowheads. Magnified segments on the left are used to show the different spine subtypes (D2): mushroom (m), stubby (s) and thin (t). **E:** Graphs showing changes in total density of dendritic spines corresponding to the first 150μm from soma and in distinct segments stablished (0-50μm, 50-100μm and 100-150μm). **F-H:** Graphs showing total spine density considering whole length (150µm) and in specific segments of 50µm for mushroom (F), stubby (G) and thin spines (H). **I-J:** Structural Sholl analysis of GAD-EGFP and PV expressing interneurons in the *stratum oriens*. The images show 2D projection of a Z confocal stack of GAD-EGFP (I) and PV (J) positive interneurons. **I1-2 and J1-2**: Representative 3D reconstructions of dendritic arbors from rhEPO and PL treated mice. Graphs show changes in total number of intersections with Sholl spheres and in number of intersections as function of distance from soma. All graphs show mean±SEM; N numbers depicted as dots in the bars; unpaired two-tailed Student’s *t*-test. All analyses were conducted following treatment scheme in B, at same age, and same area evaluated. Scale bar: 5μm for C and D; 6.5μm for 3D projections in I-J and 20μm for 3D reconstructions in I-J.

To examine whether rhEPO induces changes in the synaptic output of SST O-LM cells, we first studied the density of GAD-EGFP positive *en passant* boutons (*EPB*) in the *stratum lacunosum-moleculare* (**Figure 4C**), where the axons of SST O-LM cells ramify and establish synaptic contacts with the distal dendritic tuft of pyramidal cells and other interneurons. After rhEPO, we noticed a slight decrease in *EPB* density (*t*=1.955; *p*=0.0791), suggesting a reduced neurotransmitter release and thus higher excitability of pyramidal neurons.

Due to the peculiarity of SST O-LM displaying dendritic spines along their dendrites, we analyzed the dendritic spine density in GAD-EGFP+ interneurons^67^ (**Figure 4D**). When considering the whole length of dendrites, rhEPO reduced the total density of dendritic spines (**Figure 4E**; *t*=2.565; *p*=0.0281). This was primarily owed to the reduction of dendritic spine density in the most distal segment, i.e. from 100 to 150μm (*t*=3.991; *p*=0.0026). We next analyzed the morphology of dendritic spines, taking into account three distinct spine subtypes (**Figure 4F-H**), namely mushroom, stubby and thin. After rhEPO, the total density of mushroom spines was markedly decreased (**Figure 4F**; *t*=3.055; *p*=0.0121), specifically in the more distal segments: 50-100μm (*t*=2.024; *p*=0.0705) and 100-150μm (*t*=3.085; *p*=0.0115). This subtype corresponds to the most mature and stable type of dendritic spines. Additionally, while we noticed a reduction in stubby spine density in the most distal segment (**Figure 4G**; *t*=2.914; *p*=0.0154), the density of thin spines remained unaltered.

Finally, we quantified the complexity of the dendritic arbors of SST O-LM and PVALB interneurons (PV+) using Sholl analysis. The total number of dendritic intersections of SST O-LM, identified as GAD-EGFP+ cells, tended to decrease upon rhEPO (**Figure 4I**; *t*=2.197; *p*=0.0527). Considering the number of intersections in the different Sholl spheres, we found a reduction in the first 80μm of dendrite length: 40μm-radius (*t*=2.595; *p*=0.0267), 60μm-radius (*t*=2.580; *p*=0.0274) and 80μm-radius (*t*=3.131; *p*=0.0107). Sholl analysis in PV+ interneurons also revealed a reduction in the total number of intersections (**Figure 4J**; *t*=2.458; *p*=0.0338) and specific Sholl spheres: 50μm-radius (*t*=1.848; *p*=0.0977), 60μm-radius (*t*=2.836; *p*=0.0195), 70μm-radius (*t*=2.317; *p*=0.0457) and 80μm-radius (*t*=2.012; *p*=0.0751). Taken together, we demonstrate that structural complexity and neurotransmitter release capacity of certain interneuronal subpopulations are compromised upon rhEPO, suggesting less inhibition of pyramidal neuronal excitability in hippocampus.

### rhEPO treatment marginally affects excitatory/inhibitory (E:I) balance in the hippocampal parenchyma

We subsequently investigated the impact of rhEPO on E:I balance in the hippocampus. As the phenomenon of brain homeostasis might compensate for such imbalance, we did not expect a dramatic E:I disturbance in whole parenchyma upon rhEPO, but a noticeable difference. We asked whether rhEPO may affect the expression of excitatory and inhibitory presynaptic markers (**Figure 5A**) and thus analyzed the density of puncta expressing VGLUT1 and VGAT as a proxy for synaptic density, reflected as E:I ratio (**Figure 5B-E**). While our results show, in general, a trend of higher E/I ratio after rhEPO, expression of these excitatory and inhibitory markers was not significantly different. Nonetheless, we observed a slight increase in VGAT+ puncta density in *stratum radiatum* (**Figure 5D**; *t*=2.178; *p*=0.0574), leading to reduction of E/I balance (density of VGLUT1+ puncta/density of VGAT puncta) (*t*=2.205; *p*=0.0549). Collectively, we add another layer to our above findings: EPO seems to reconfigure synapses, probably by fine-tuning the balance between excitatory and inhibitory neurons.

**Figure 5.**
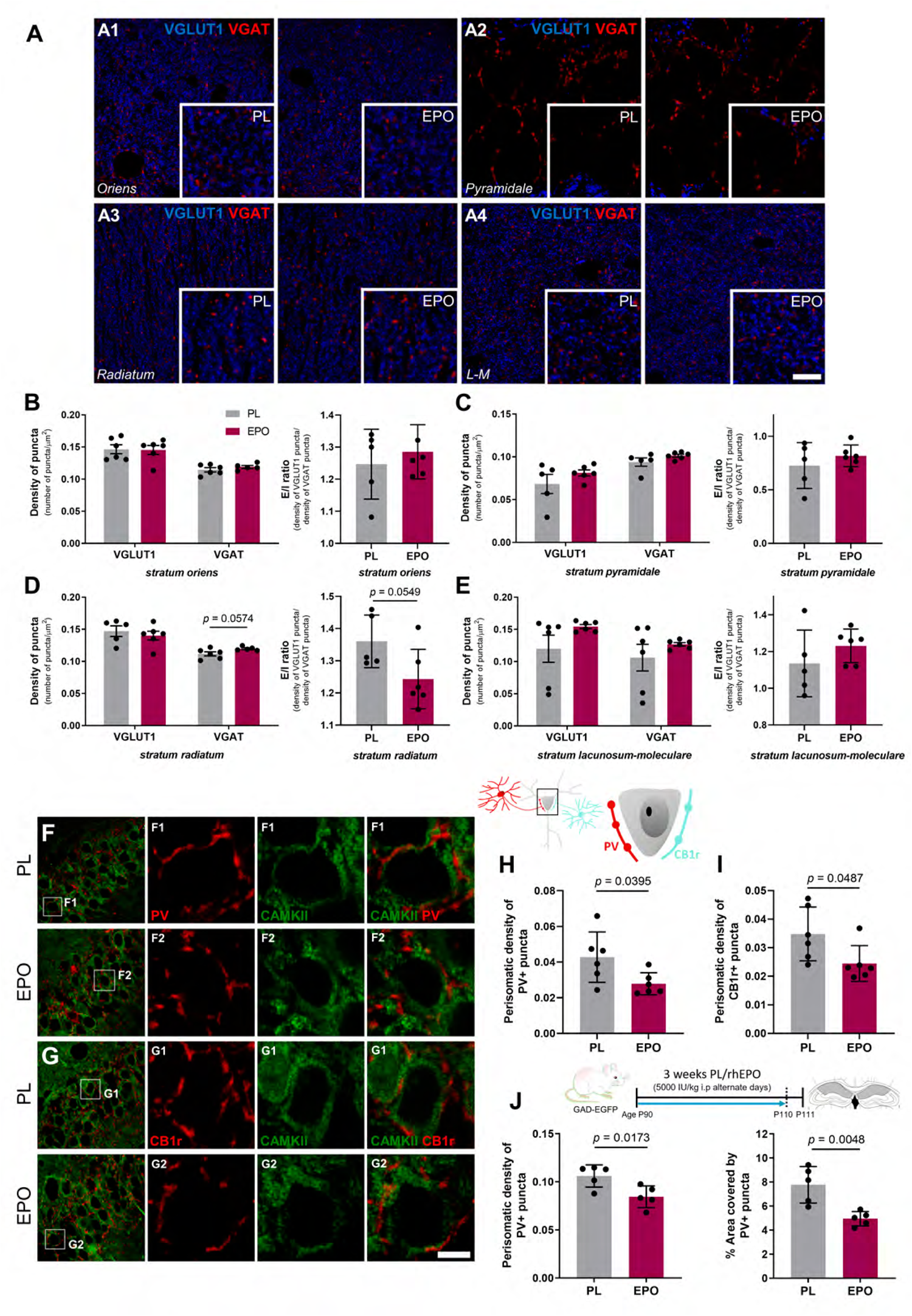
Treatment with rhEPO slightly affects the E:I ratio and decreases the density of inhibitory perisomatic puncta on excitatory hippocampal neurons. **A:** Analysis of excitatory/inhibitory balance in CA1. Single confocal planes with magnified insets show expression of VGLUT1 (blue) and VGAT (red) immunoreactive puncta from rhEPO and PL treated mice in the CA1 strata: *oriens* (A1), *pyramidale* (A2), *radiatum* (A3) and *lacunosum-moleculare* (L-M; A4). **B-E:** Graphs showing density of puncta expressed as number of puncta *per* micron and E/I ratio in the different layers. **F-I:** Analysis of the density of perisomatic inhibitory puncta on excitatory neurons. Schematic illustration shows perisomatic input that pyramidal neurons receive from both types of basket cells, PVALB and CCK basket cells (PV+ and CB1r+ puncta, respectively). **F-G:** Panoramic and single confocal views of CA1 *stratum pyramidale* showing PV and CB1r expressing puncta (red), surrounding the soma of CAMKII+ excitatory neurons (green). **H-I:** Graphs present changes in density of perisomatic puncta expressing PV (H) and CB1r (I). **J:** Density of perisomatic inhibitory puncta in CA1 of older GAD-EGFP mice. Graphs show changes in density of perisomatic PV+ puncta and in percentage of area covered by PV expressing puncta. All graphs show mean±SEM; N numbers depicted as dots in the bars; unpaired two-tailed Student’s *t*-test. All analyses were conducted following treatment scheme in Figure 4B, except Figure 5F that followed the 3-week PL/rhEPO treatment at older age (P90). Scale bar: 5μm for A; 25μm for F-G panoramic views and 5μm for insets.

### rhEPO treatment decreases the density of inhibitory perisomatic puncta on excitatory neurons

Among the wide variety of interneuronal lineages, basket cells are the most abundant in the favorite strategic position of shaping synaptic properties of their target neurons. Thus, we analyzed the communication between pyramidal neurons and basket cells upon rhEPO, by quantifying the perisomatic innervation that CAMKII-labelled pyramidal neurons receive from PVALB and CCK basket cells, the latter characterized by presynaptic expression of cannabinoid 1 receptor (CB1r). We determined the perisomatic density of PV and CB1r puncta around individual principal cells in *stratum pyramidale* (**Figure 5F-I**). In adolescent mice (P49), we observed a decrease in PV+ and CB1r+ puncta density after rhEPO (**Figures 5H-I**; PV+ puncta: *t*=2.367; *p*=0.0395. CB1r+ puncta: *t*=2.243; *p*=0.0487). Similarly, in adult mice (P111; **Figure 5J**) rhEPO decreased density (*t*=2.990; *p*=0.0173) and percentage of area covered by PV+ puncta (*t*=3.860; *p*=0.0048). Altogether, rhEPO attenuates the cross-talk of interneurons with mature pyramidal neurons by influencing basket cells from both PVALB and CCK subsets.

### Electrophysiological recordings elucidate a shift in the hippocampal excitatory/inhibitory balance (E:I) upon rhEPO

We next explored functional facets of rhEPO treatment on hippocampal activity (**Figure 6A**). In a virtual reality environment, we obtained in awake mice local field potential (LFP) recordings of the pyramidal layer of the dorsal CA1 subfield. With this design, we induced exploratory behaviors in the animals, a condition evoking an active hippocampal state in form of theta rhythmicity.

**Figure 6.**
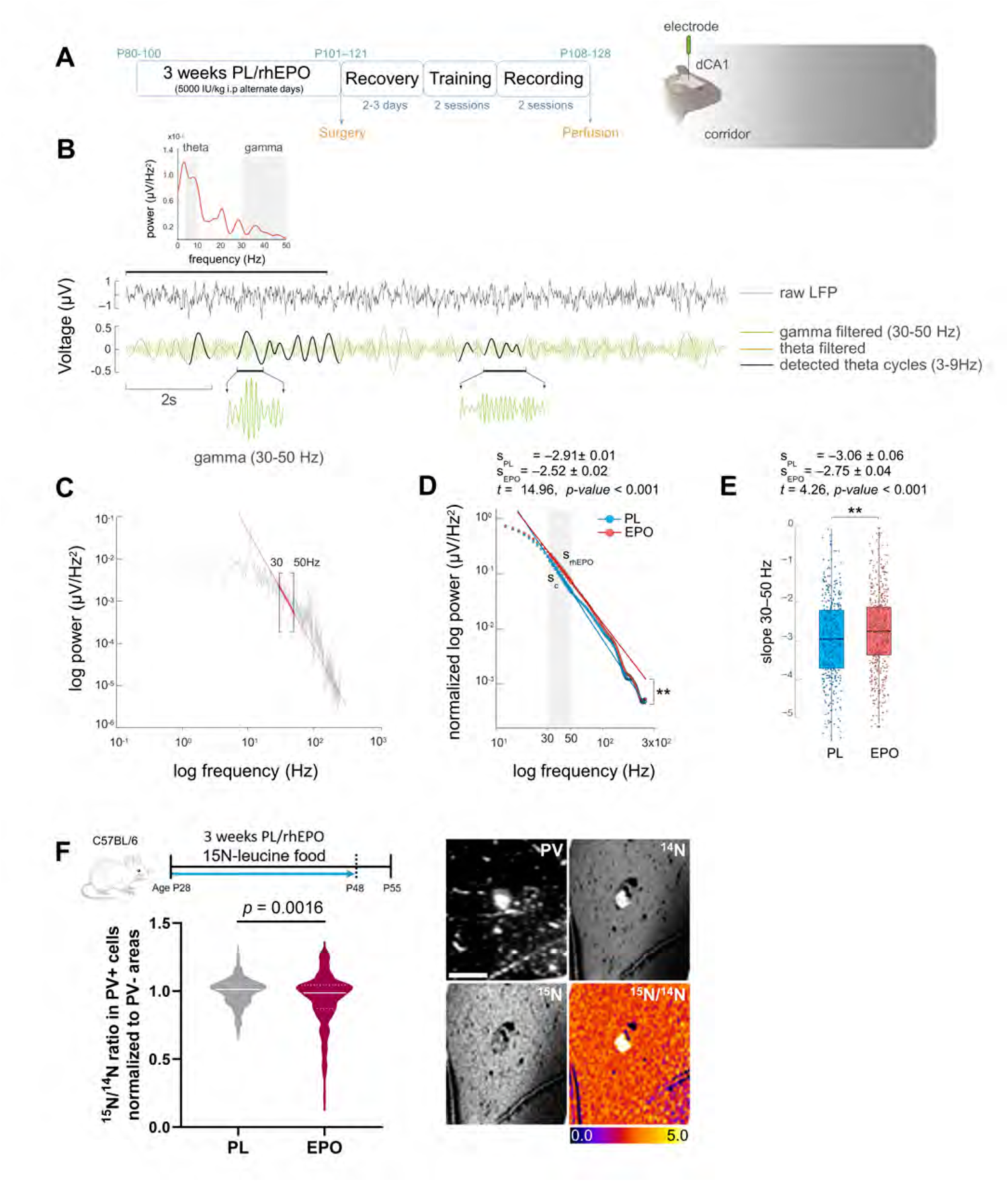
rhEPO produces a shift towards excitation and decreases the metabolic activity of hippocampal interneurons. **A:** Scheme of experimental timeline and diagram of recording electrode location in the dorsal region of CA1 (dCA1). **B:** Representative power spectrum and raw LFP signal obtained during recording. After filtering, theta cycles and their coupled gamma activity are highlighted with black and green lines, respectively. **C:** Power spectrum highlighting the mean slope of the gamma activity of interest (30-50Hz) with a red line. **D:** Non-linear regression model of normalized power spectrum showing a decrease in the mean slope of the 30-50Hz activity band of rhEPO treated mice. **E:** Boxplot of slope values obtained in the 30-50Hz band showing a decrease in the slope of rhEPO treated mice. **F:** Schematic of NanoSIMS experiment after ^15^N-leucine-enriched food, provided for the same 3-week duration as PL/rhEPO treatment (N=4 mice/group), starting at P28. Representative images and calculated ^15^N/^14^N ratio in PV+ cells as measure of ^15^N-leucine incorporation (normalized to PV-areas). Electrophysiological statistical analysis explained in text; for NanoSIMS: 2-tailed Mann-Whitney *U* test; scale bar: 15μm for F.

Selecting theta epochs on hippocampal LFP, we filtered the raw signal to detect presence of coupled gamma waves (**Figure 6B**), a typical oscillatory profile of hippocampal theta activity, reflecting periodic bouts of excitation and inhibition^68^.

According to Gao and colleagues^69^, we explored a slope-fitting model of the power spectra on the 30-50Hz band to capture differences in E:I ratio (**Figure 6C**). We found a decrease of the 30-50Hz slopes in rhEPO treated mice (placebo: s_PL_=–2.91; rhEPO: s_EPO_=–2.52; *t*=14.96, *p*<0.001; **Figure 6D**), calculated by regression models on the mean power spectral density (PSD) for both groups. We confirmed this result by comparing the means of 30-50Hz slopes for both groups, calculated from the PSD of each theta cycle (placebo: s_PL_=–3.06; rhEPO: s_EPO_=–2.75; *t*=4.26, *p*<0.001; **Figure 6E**).

### rhEPO decreases the metabolic activity of hippocampal PV-expressing cells

We used nanoscale secondary ion mass spectrometry (NanoSIMS)^19,20,70^ for analyzing the metabolic turnover in the CA1 PVALB subpopulation (PV+) through ^15^N-leucine incorporation (**Figure 6F**). This method provides indirect evidence of protein synthesis, commonly associated with e.g. growth processes or cellular activity. When we compared the ^15^N/^14^N ratio in PV+ cells, we observed a significant decrease after rhEPO (*p*=0.0016), indicative of an abridged metabolic turnover. This contrasts to the increased ^15^N-leucine incorporation previously observed upon rhEPO in NeuN+ cells of the *stratum pyramidal*e^19^. In parallel to the rhEPO stimulated neurodifferentiation of pyramidal cells in CA1, resulting in substantial numbers of new functional neurons with increased metabolism and density of dendritic spines^17,19^, we observed a decrease in structural complexity, connectivity, and metabolism in specific interneuronal subpopulations.

### rhEPO alters the expression of plasticity-related molecules associated with hippocampal interneurons

Certain types of plasticity-related molecules play key roles in the structural remodeling and connectivity of neurons, specifically interneurons^31,32,71,72^. We initially performed a densitometric analysis of polySia-NCAM expression in the different layers of the CA1 region (**Figure 7A**). Increases after rhEPO treatment were observed in the *strata oriens* (*t*=2.602; *p*=0.0315), *radiatum* (*t*=2.782; *p*=0.0239) and *lacunosum-moleculare* (*t*=3.076; *p*=0.0152; **Figure 7B**). A tendency towards an increase was also observed in the *stratum pyramidale* (*t*=1.078; *p*=0.0833). Since most PNNs in the hippocampus surround PVALB cells, we also analyzed the total number of PV+ cells, PNNs and their co-localization in the CA1 region (**Figure 7C**). Animals treated with rhEPO displayed an increased number of PV+ (*t*=3.334; *p*=0.0103) and PV+PNN+ (**Figure 7D**; *t*=2.963; *p*=0.0181) cells. The number of PNNs also tended to be elevated (*t*=2.171; *p*=0.0617) after rhEPO. In agreement with earlier results in humans and mice^15,19,73^, volumetric analysis of CA1 area revealed a tendency of an increase upon rhEPO (**Figure 7E**; *t*=1.203; *p*=0.1316).

**Figure 7.**
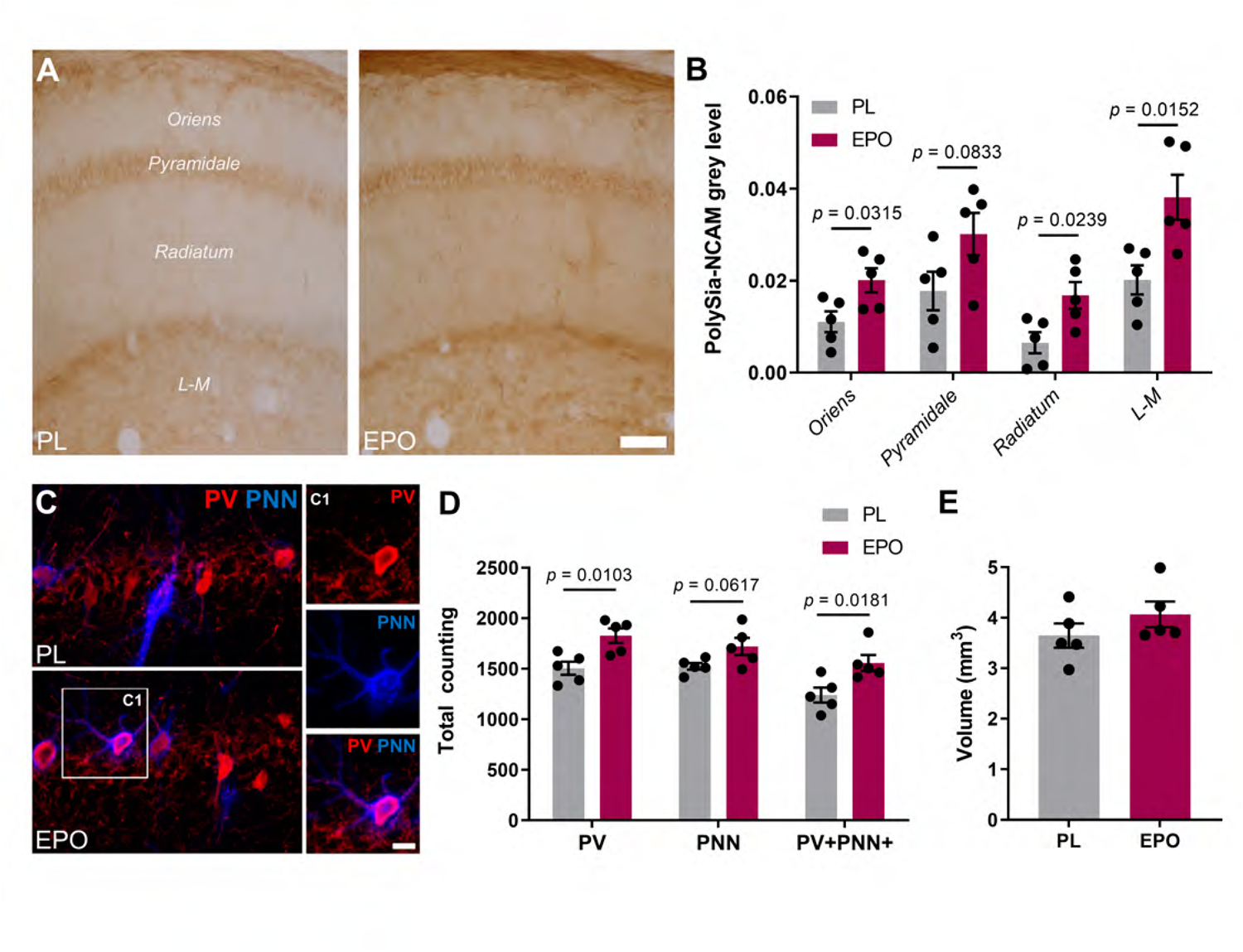
Treatment with rhEPO decreases the expression of plasticity-related molecules in the hippocampus. **A:** Densitometric analysis of polySia-NCAM expression in CA1. Microphotographs from conventional light microscope compare the expression of polySia-NCAM in the CA1 strata (*oriens, pyramidale, radiatum* and *lacunosum-moleculare*) in rhEPO and PL treated mice. **B:** Graph representing changes in the grey levels of polySia-NCAM immunoreactivity. **C:** Expression of PV, PNNs and their co-localization after rhEPO treatment. Confocal plane focused on *stratum pyramidale* shows distribution of PV+ interneurons (red) and PNNs (blue) in rhEPO and PL groups. The squared area (C1) shows a single PV immunoreactive neuron surrounded by a PNN. **D:** Graph presenting changes in total number of PV+ cells and PNNs, and the total number of PV+ neurons surrounded by PNNs. **E:** Graph showing the volume of CA1 in rhEPO and PL treated mice. All graphs show mean±SEM; N numbers depicted as dots in the bars; unpaired two-tailed Student’s *t*-test (except 7E, one-tailed). All analyses were conducted following treatment scheme in Figure 4B, at same age, and same area evaluated; scale bar: 60μm for A; 12.5μm for overview in C and 9μm for inset magnifications.

## DISCUSSION

Improvement of mood and higher cognition following rhEPO treatment can in principle be an indirect effect due to increase in hematocrit and thus oxygen delivery. Although we do not discard this possibility entirely, our previous findings support a hematopoiesis-independent, specific mode of rhEPO action in brain. For instance, elimination of EPOR from pyramidal neurons prevented rhEPO-induced hippocampal and behavioral changes^17^. Transgenic overexpression of spontaneously active EPOR in hippocampal pyramidal neurons enhanced higher cognition^18^. Moreover, we reported very recently that rhEPO markedly influences the transcriptome of excitatory hippocampal neurons, thereby inducing e.g. associated signaling cascades to tackle metabolic challenges^21^.

Despite the complex nature of synapses, it takes both pyramidal cells and interneurons to form viable circuits. In the present work, we identified interneuron lineages that exhibit core functional differences upon rhEPO, thus taming synapses. Employing the relevant computational approaches, we here provide a detailed transcriptomic analysis of hippocampal interneurons in mice, treated with either rhEPO or PL. Although most lineages classified in this study are consistent with previously reported interneuronal subpopulations, we found two distinct lineages that are not documented to date. These lineages exhibit distinct transcriptomes, co-express *Gad1* and *Gad2* genes at high level, but not markers of other lineages, supporting their interneuronal origin. We named them according to the top marker genes they express, *Zbtb20/Mgat4c* and *Nrg1/Ptprd*, respectively.

Upon rhEPO, we observe a substantial difference in the gene expression profiles of distinct subgroups of interneurons. To determine whether the changes in transcriptional readouts are secondary events or whether rhEPO is directly involved in modulating them, we asked if these subgroups of interneurons do express *Epor*. Indeed, we found that a generous portion of interneurons is expressing *Epor*, enough to be detected in our FISH experiments. These results affirm that the interneuron population is an addition to the known arena of EPO action, which are excitatory neurons and glial cells^17,19–21^. Therefore, this premise is pleading for further investigations.

EPOR expression by interneurons might indicate that rhEPO impacts their transcriptome, but it remains open which genes and/or transcriptional networks, *per se*, are influenced by the EPO-EPOR cascade. Our DEG analysis shows that rhEPO modulates cell-type specific transcriptomes. The ability of rhEPO to remodel the regulatory networks of host cells was recently defined in pyramidal lineages^21^. The present work extends further, demonstrating that rhEPO remodels the network of both neuronal populations with opposing characteristics. In previous studies, rhEPO was found to accelerate firing and metabolism, increase the number of neuronal progenitors, and skew the developmental trajectory of pyramidal neurons^17,19,21^.

This supports our findings that several genes associated with the functional integrity of structure and synaptic connectivity are responsive to rhEPO treatment. The exact mechanism by which rhEPO modulates synaptic architecture and structural integrity of interneurons, however, remains still unclear. Upon rhEPO treatment, we identified a distinct set of ligands expressed or up/downregulated in the interneuron subpopulation, with known receptors in the pyramidal subpopulations. The secreted SLIT protein families, *Cadm1*, and neuroligins (*Nlgn1/2/3*) expressed from interneuron lineages binding to ROBO receptor, *Nectin3*, and neurexins (*Nrxn1/2/3*), respectively, on pyramidal lineages are among the most abundant interactions in our datasets. In agreement with the consensus of ligand-receptor dynamics^59,74,75^, *Nrg3* signal from PVALB axo-axonic is received by *Erbb3/4*^60^, however, we find here that *Nrg3* also interacts with *Egfr* that is expressed by pyramidal neurons. While these housekeeping interactions show no difference between rhEPO and PL subjects, there were a few interactions distorted in rhEPO samples. For instance, *Calm2-Pde1c* and *Pdgfc-Pdgfrb* pairs were highlighted in this study where the former interaction seems to have been abolished and the latter was built in rhEPO samples. These interactions exhibit a broad spectrum of functions in brain and have been shown to regulate neurogenesis, cell survival, and synaptogenesis in hippocampus^65,76–80^.

We note that our analysis is limited by the in-silico analysis of receptor-ligand interactions and many possible confounders such as spatial and temporal states of these cells in the hippocampus which we could not control for. Yet the shifts in receptor expression we observed are intriguing and merit further investigation in future studies.

Our *in/ex vivo* studies revealed rhEPO as a potent factor influencing the structural and synaptic plasticity of certain inhibitory neurons. Following 3-week rhEPO treatment, we found the complexity of dendritic arbors and density of mature dendritic spines of SST O-LM cells markedly reduced. The opposite effects that rhEPO administration has on CA1 pyramidal neurons, namely an increase in their dendritic spine density and in the number of mature neurons^17,19^, might enhance the excitatory input that SST O-LM cells receive from pyramidal neurons and prompt them to respond in a protective manner by reducing their dendritic features. We previously noticed similar dynamics in spiny SST-expressing interneurons within the basolateral amygdala after chronic stress. While this paradigm increases the dendritic complexity and the spine density of excitatory neurons, SST interneurons face a noticeable atrophy^81^. This aligns also well with our previous studies where glutamate, acting through NMDA receptors, has a profound effect on the structure and connectivity of SST O-LM cells^36,82^, suggesting that the rhEPO-induced changes in glutamatergic neurons may cause the markdown of dendritic arborization and spine density observed in this subpopulation. This would make SST cells less inhibitory, which is reflected in the reduction of their axonal *en passant boutons* in the *stratum lacunosum-moleculare,* the main output of this subpopulation. We observe similar dynamics in PVALB interneurons of the *stratum oriens*, where their dendritic complexity is compromised, possibly in response to the enhanced excitatory input^17^. This dampening at the structural level is accompanied, as in SST neurons, by a reduction in their synaptic output onto pyramidal neurons, as demonstrated by the decrease in density of PV+ perisomatic puncta around pyramidal neurons, and in a reduced metabolic protein turnover of PVALB interneurons. The reported impairment of gamma oscillations in mice with selective activation of EPOR signaling in GABAergic neurons^83^ is consistent with these findings, since PVALB basket cells regulate gamma oscillations in CA1 through their perisomatic innervation of pyramidal neurons^84^. Similarly, we report here a reduction in the density of CB1r+ perisomatic puncta, indicating that CCK basket cells, which also regulate hippocampal gamma oscillations^85^, may be another target of rhEPO, responding alike PVALB interneurons. We show here that - similar to the pyramidal lineage^17^ - these changes in the interneuronal circuitry are maintained from adolescence to adulthood, suggesting long-term improvement of cognition by the double-edged effect of rhEPO.

The observed synaptic renovation by rhEPO could largely be explained by reduced firing rates in inhibitory and/or accelerated firing in excitatory neurons enhancing the overall excitability. However, it is also possible that the response of inhibitory neurons precedes that of pyramidal neurons upon rhEPO^17^. Moreover, compensatory phenomenona may help maintain homeostasis, including connectivity with interneuronal subpopulations, like chandelier or interneuron-specific (IS) cells, or even variation in excitatory input coming from extrahippocampal sources.

Additionally, we show the effect of rhEPO on plasticity-related molecules such as polySia-NCAM and PNNs, associated with hippocampal inhibitory interneurons. The expression of polySia-NCAM strongly regulates structure and connectivity of hippocampal interneurons, including SST O-LM and PVALB basket cells^37,86^. The presence of polySia-NCAM, which is enhanced after rhEPO treatment, is associated with reduced dendritic arborization, density of spines and synaptic contacts^67^, while its depletion rescues them all^37,86^. We also notice an increase in PNNs, matricellular structures that mainly enwrap PVALB interneurons^27,30,32^. PNNs could promote synapse stabilization and restrict plasticity processes^87^, suggesting a role in the stabilization of hippocampal circuits, which complies with EPO-induced improvement of higher cognition^12,19^. We likewise observed an increase in the total number of PVALB cells, suggesting a shift in the PVALB network configuration towards being more prone to cognitive enhancement and structural synaptic plasticity in the hippocampus^88^. Thus, changes in interneuronal structure, connectivity and physiology may have been mediated, at least partially, by rhEPO via interneuronal plasticity-related molecules.

Our results seem to somewhat contradict those of Khalid and colleagues who reported accelerated maturation of GABAergic neurons and enhanced inhibitory neurotransmission upon rhEPO. These discrepancies can be due to the type of model used, i.e. Tg21 mice, that constitutively overexpress EPO in brain during their whole life, different methodological approaches and time points investigated^89^.

In summary, while EPO drives a series of transcriptomic and physiological changes in diverse interneuronal populations, our final aim is still to identify the precise mechanism underlying the observed benevolent phenotypes. Nonetheless, this study sets a reliable foundation to investigate the cell-level communication of various lineages of interneuronal and pyramidal layers that help to alleviate mood, memory and cognition-associated disorders. The impact of rhEPO treatment on interneurons, especially on SST and PVALB cells may have important clinical implications, since alterations in these inhibitory cells have been described in different psychiatric disorders^90^. Moreover, aversive experiences, especially during early life, which are known to be predisposing factors for these disorders, also have profound effects on the structure and connectivity of interneurons^31,91–93^. Consequently, new therapeutic approaches, such as rhEPO treatment, able to modulate inhibitory neurons as well as the expression of molecules related to their plasticity, may be promising candidates for innovative strategies.

## EXPERIMENTAL PROCEDURES

### Experimental Models

Twelve juvenile transgenic male mice [GIN (GFP-expressing Inhibitory Neurons), Tg(GadGFP)45704Swn] from Jackson Laboratories (Bar Harbor, Maine, United States) were used at the starting age of P28 for the rhEPO treatment study. Under the control of the mouse *Gad1* gene promoter, these mice express the enhanced green fluorescent protein in a subpopulation of SST-expressing interneurons, GAD-EGFP^94^. Due to the dimensions of the skull in the electrophysiological study (only possible in adult mice, P80-P100), an additional cohort of GAD-EGFP mice (P90, n=10) was incorporated in this study to validate some results at this older age. Different sets of C57BL/6 mice (Charles River) were used for the extracellular recordings (P90, n=10), the snRNA-sequencing analysis (snRNAseq; P28, n=23) and the L-leucine incorporation study (NanoSIMS experiment; P28, n=8). Three extra mice at the age of P90 were used for characterization of the EPOR expression. All mice were housed in a temperature-controlled environment (21 ± 2 °C) on a 12 h light–dark cycle with food and water available *ad libitum*. All treatments and methods were carried out following the animal care guidelines of the European Commission 134 2010/63/UE and the experimental design was conducted in accordance with the ethics committee of the University of Valencia (2020/VSC/PEA/0106) and the local Animal Care and Use Committee (Niedersächsisches Landesamt für Verbraucherschutz und Lebensmittelsicherheit, LAVES – AZ 33.19-42502-04-17/2393) following the German Animal Protection Law. Every effort was made to minimize the number of animals used and their suffering.

### Mouse Treatment

Recombinant human (rh)EPO (5000 IU/kg body weight; NeoRecormon, Roche, Welwyn Garden City, UK) or placebo (PL; solvent control solution EPREX®buffer) were applied via i.p. injections (0.01 ml/g) every other day for 3 consecutive weeks with a total of 11 injections. Gin male mice at the age of P28 or P90 (juvenile vs adult studies) followed the 3-weeks PL/rhEPO paradigm and were sacrificed 24h after the last injection (P49 and P111, respectively). C57BL/6 mice for snRNAseq study underwent the same procedure at the age of P28. For nanoscale secondary ion mass spectrometry (NanoSIMS) study, mice obtained food pellets containing 1.025% L-leucine-15N stable isotope (Sigma-Aldrich, St Louis, MN, USA) for 3 weeks between P28 and P49 (in parallel with PL/rhEPO injections). Based on previous results^17,20^, mice were sacrificed after 1 week break (P55). C57BL/6 mice used for electrophysiological studies initiated the 3-weeks PL/rhEPO treatment between P80 and P100. Mice were weighted before each injection to receive the appropriate dose. During all the procedures, the experimenter was blind to group assignment.

### Perfusion and Microtomy

Mice were deeply anesthetized with sodium pentobarbital (150 mg/kg) and transcardially perfused via left cardiac ventricle with saline (NaCl 0.9%) or Ringer’s solution followed by 4% paraformaldehyde (PFA) in sodium phosphate buffered saline (PBS) 0.1 M, pH 7.4 solution. The right hemisphere, destined to neuronal structural analysis, was cut in 100-μm-thick coronal sections with a vibratome (Leica VT 1000E, Leica), collected in 3 subseries and stored at 4°C in PB 0.1 M with sodium azide (0.05%). The left hemisphere, intended for immunohistochemical analyses, was cryoprotected with 30% sucrose in PBS 0.1 M for 48 h and cut afterwards in 50-µm-thick coronal sections using a sliding microtome (LEICA SM2000R, Leica). Slices were collected in 6 subseries and stored at −20°C in a cryoprotective solution (30% glycerol, 30% ethylene glycol in PBS 0.1 M). Brains destined to NanoSIMS analysis were processed as before but coronal sections of 30 µm thickness were obtained with a Leica CM1950 cryostat (Leica Microsystems, Wetzlar, Germany) and stored at −20°C in 25% ethylene glycol and 25% glycerol in PBS.

### Immunohistochemistry

For fluorescence immunohistochemistry, “free-floating” sections were blocked and permeabilized with 5% normal donkey serum (NDS, Jackson ImmunoResearch Laboratories) in 0.3% Triton X-100 in PBS (PBST; Sigma-Aldrich) for 1h at room temperature (RT). Subsequently, slices were incubated for 48 h at 4 °C with the appropriate primary antibody cocktail diluted in 5% NDS with 0.3% PBST (**Table S4**). After washing, sections were incubated for 2 h at room temperature with different secondary antibody cocktails (**Table S4**) diluted in 3% NDS with 0.3% PBST. Finally, sections were washed in PBS 0.1 M, mounted on slides and coverslipped using Dako Fluorescent Mounting Medium (Dako Diagnósticos). Sections were processed together for immunohistochemistry to reduce the variability of staining and allow comparisons between groups.

To analyze the expression of polySia-NCAM in conventional light microscopy, sections were processed using the ABC method. After several washes, sections were first incubated for 1 min in an antigen unmasking solution (0.01 M citrate buffer, pH 6) at 100 °C. After cooling down the sections to RT, they were incubated with 10% methanol and 3% H_2_O_2_ in PBS for 10 min to block endogenous peroxidase activity. After this, sections were blocked and incubated with the primary antibody as mentioned before. After washing, sections were incubated for 2 h (at RT) with a secondary biotinylated antibody followed by an avidin-biotin-peroxidase complex (ABC; Vector Laboratories, Peterborough, UK) for 1 hour in PBS. See Table S4 for antibody information. Color development was achieved by incubating with 0,05% 3,3′-diamino-benzidine tetrahydrochloride (DAB; Sigma–Aldrich) and 0.033% H2O2 for 4 min. Finally, sections were mounted on slides, dried for 1 day at RT, dehydrated with ascending alcohols and rinsed in xylene. After this, sections were coverslipped using Eukitt mounting medium (PANREAC).

### Surgery

At 24 hours after the last PL/rhEPO injection, mice were anesthetized with isoflurane (4% for induction, 2% for maintenance, both in 0.5 ml O_2_/min flow rate). Additionally, atropine (0.1 mg/kg) and buprenorphine (0.1 mg/kg) were s.c. injected to minimize respiratory pitfalls and suffering during surgery. Ophthalmic ointment was applied to protect the eyes during surgery. Animals were placed in a stereotaxic frame (Narishige), then lidocaine (2%, Normon) was s.c. injected above the incision area and the skull surface was exposed and cleaned with 70% ethanol. Afterwards, the skull was trephined in two locations in the following coordinates relative to bregma: AP - 2.00, ML −1.50 for the subsequent *in vivo* recording of the right dorsal hippocampus, and AP +0.30, ML +0.50 to place a reference electrode (formvar insulated stainless steel monopolar macroelectrode (120 μm in diameter, WPI) in the subarachnoid space. Then, a polymeric resin (OptiBond; KERR) was applied on the skull surface and polymerised using a UV light lamp (Woodpecker Led D). Afterwards, a custom-made 3D printed resin headplate was placed on the skull and dental cement (DuraLay, Reliance Dental) was applied to cover all the tissue exposed except the cranial window, sealed with SILASTIC (Kwik-cast, WPI). After surgery, mice were s.c. injected with buprenorphine (0.1 mg/kg) and meloxicam (3 mg/Kg) and placed under a heat lamp and monitored until complete recovery.

### Electrophysiological Recording

#### Training

At 48 hours after surgery, mice were trained for two consecutive days to get familiarized with our custom-made virtual reality platform (CIBERTEC, Spain). After a water restriction period, ranging randomly from 1 to 3 hours, mice were placed in a head-fixed frame on a low-friction styrofoam cylinder and positioned in front of a TFT curved screen. To simulate virtual navigation, cylinder movement was recorded using an infrared sensor applied on an open-source electronic platform (Arduino Mega board, 2560 Rev3) and converted into movements of custom-made virtual corridors implemented in Unity3D software (Unity Technologies, 2018). Four corridors (C1-C4) were configured and designed with colours within the range of wavelengths perceived by the mouse visual system^95,96^. The virtual corridors were split into four virtual sectors (S1-S4) based on distinct cue patterns on the walls. A distinctive cue (white cross) identified a final reward area, where a 1% sucrose dilution on water drop was provided. Animals were left undisturbed to explore twice the same corridor without time limitation.

#### Recording

Twenty-four hours after the end of training, electrophysiological recording of the mice was carried out in one session with full corridor exploration. After placing them in the head fixation frame, a cranial window was exposed, and the local field potential (LFP) of the dorsal hippocampus (CA1) was recorded with a 120 μm diameter Teflon-coated steel monopolar macroelectrode (WPI), whose tip was located 1.30 mm ventral to the brain surface. The signal was referenced against the electrode previously placed in epidural space and acquired using the Open-ephys system^97^ with a sampling frequency of 30 kHz. Then, raw signals were then imported to the Matlab development environment (The MathWorks, Natick, MA, USA) and analyzed using built-in self-developed routines or standard MATLAB libraries as necessary.

#### Data analysis and statistics

The hippocampal recording was incorporated into the MATLAB framework as a time series of the LFP measures. First, the recordings were downsampled to 1000 Hz, digitally notch-filtered with a Butterworth bandstop filter of around 50 Hz (and its harmonics) and z-scored to normalize the original signals.

It is well-known that during active exploration, theta (5–12 Hz) oscillations dominate the hippocampal CA1 area of the rodent brain^98^. We extracted the theta periods by decomposing the normalized signal into the different predominant frequency bands by the empirical mode decomposition (EMD, Hilbert-Huang transform), thereby preserving the time domain during periods of active exploratory behavior (animal velocity > 2 cm/s). We isolated the low-frequency (< 5Hz), theta (5-12 Hz) and supra-theta (> 12 Hz) signals by combining the components with mean instantaneous frequencies into the ranges < 5 Hz, 5-12 Hz and >12 Hz, respectively.

We then identified individual theta cycles by detecting each cycle’s local maxima and minima with an absolute value above the envelope of the low-frequency time series. A theta cycle was thus defined by a candidate central peak surrounded by a pair of consecutive valleys, separated at least by 71 ms (∼14 Hz) and no more than 200 ms (∼5 Hz).

The power spectral density (PSD) from all theta cycles was estimated by computing the Fourier transform (Welch method with 50% overlapping and a Hamming 5 s window at a resolution of 0.2 Hz). Using the log-log representation of the PSD, we used linear regression to find the slope of the best fit over the 30–50 Hz frequency range^69^.

Statistical analysis was performed in Rstudio (4.2.3). Since we worked with the slope values derived from a regression model on the PSD data in the frequency range 30-50 Hz, we first aimed to apply a nonlinear regression fit on the mean PSDs of each experimental group:

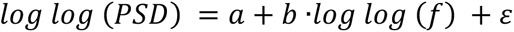

In addition, as a second approach, we wanted to apply inferential statistics on the values of the 30-50 slopes of each theta cycle extracted from the original signals. In this case, the applicability conditions of parametric tests were evaluated with the Shapiro-Wilks normality test. Finally, having different samples for both groups, Welch’s test with a significance level of p < 0.05 was used to determine significant differences. All results were expressed as mean ± SEM. Graphs were obtained using the R package ggplot2. Data access (DOI: 10.5281/zenodo.7885936).

### Structural Analysis of GAD-EGFP and PV Expressing Interneurons

All the structural parameters of GAD-EGFP+ (∼SST O-LM) and PVALB interneurons were studied using a laser scanning confocal microscope (Leica TCS SPE) as described before^67^. We focused our structural studies in the *stratum oriens* of the hippocampal CA1 region. The majority of GAD-EGFP+ interneurons located in this area can be considered SST-expressing O-LM cells, although some hippocamposeptal cells have also their somata located in this area^36^.

We initially studied the synaptic output of SST O-LM cells with somata located in the *stratum oriens* by analyzing the plexus of GAD-EGFP+ axons present in the *stratum lacunosum-moleculare* of the CA1 region. To study the density of *en passant boutons (EPB)*, we used a 63× oil immersion objective with a 2.5× digital zoom. Six axonal segments, measuring at least 10 μm, were chosen randomly *per* animal. In order to avoid an overestimation, the axonal varicosities were only considered when they fulfilled three criteria^36^: (1) they should be at least two times brighter than the axonal backbone; (2) they should be two times wider than the axonal backbone; and (3) they should not have any crossings from other axons nearby.

For the analysis of dendritic spines, 6 dendrites from 6 different GAD-EGFP expressing interneurons located in *stratum oriens* were randomly selected *per* animal. A 63× oil immersion objective and a 3.5× additional digital zoom was used to observe the first 150 µm of the dendrite in segments of 50 µm (Z-step size of 0.38 µm). The dendrites had to keep the following criteria to be included in the study: (1) their length should be at least 150 µm, and (2) no other dendrites should be found crossing their trajectory. Data were expressed as the total number of spines in the proximal (0–50 µm), medial (50–100 µm), and distal (100–150 µm) segments of the dendrite, depending on its distance from the soma. The total density of spines (density of spines in the entire length of the segment) was also analyzed. The same analysis was performed taking into account the different types of dendritic spines, classified manually according to the length of the protrusion and the diameters of their head and neck. Three different categories were established as described before in these hippocampal interneurons^37^: (1) stubby, when the length of the protrusion was <1 µm and no neck is observed; (2) mushroom, when a clear head-like structure could be observed (maximum diameter of the head was at least 1.5 times the average length of the neck) and the total length of the protrusion was <1.5 µm; and (3) thin, when the length of the protrusion was >1.5 µm, or when this length was between 1 and 1.5 µm and a clear head could not be distinguished.

For the study of the dendritic arborization, 6 GAD-EGFP- and 6 PV-expressing interneurons *per* animal were randomly selected. Z-series of optical sections (0.8 μm step size; 40× objective) covering the whole dendritic tree of selected interneurons were imaged using the sequential scanning mode. To be suitable for analysis, these interneurons had to fulfill the following features: (1) the dendritic arbor of the cell must show at least a process with a length >200 μm for the GAD-EGFP interneurons and >100 μm for the PV+ cells (due to their shorter dendrites), and (2) the soma must be located at least 30 μm deep from the surface of the tissue. The stacks obtained were then processed using FIJI/ImageJ Software^99^ to obtain 3D reconstructions. Neurons were traced using the “Simple neurite tracer” plugin, which also allowed us to analyze their Sholl profile in 3D^100^. The Sholl analysis consists on the measure of the number of intersections of the dendrites with spheres of increasing radius centered in the soma^101^. The separation among the spheres of the analysis was set at 20 μm for the GAD-EGFP interneurons and at 10 μm for the PV+ cells.

### Analysis of the Excitatory/Inhibitory Balance

We analyzed the density of neuropil puncta expressing the vesicular glutamate (VGLUT1) and GABA (VGAT) transporters, as a redoubt of the excitatory and inhibitory level, respectively, in the different layers of the hippocampal CA1 region. Confocal *z*-stacks covering the whole depth of the sections were taken with 0,38 μm step size (63x oil objective and 2x digital zoom magnification) and a laser scanning confocal microscope (Leica TCS SPE, Germany). Only confocal planes with the optimal penetration level for each antibody were selected. On these planes, 16 small squares of the neuropil (336 μm^2^) *per* layer and animal were selected for analysis to avoid blood vessels and cell somata. Images were processed using FIJI/ImageJ software as described before^102,103^: the background was subtracted with rolling value of 50, converted to 8-bit deep images and binarized using a determined threshold value. This value depended on the marker and the area analyzed and was kept the same for all images with the same marker and area. The images were then processed with a blur filter to reduce noise and to separate closely apposed puncta. Finally, the number of the resulting dots *per* layer was automatically counted and expressed as a density. The E/I ratio was calculated as the density of VGLUT1 expressing puncta divided by the density of inhibitory VGAT expressing puncta.

### Analysis of Inhibitory Perisomatic Puncta on Excitatory Neurons

To analyze the perisomatic innervation that PVALB and CCK basket cells exert, the density of puncta immunoreactive for PV or CB1r surrounding pyramidal neuron somata (identified by CAMKII expression) was analyzed as described previously^104^. Briefly, 20 CAMKII expressing neurons *per* animal located in the *stratum pyramidale* of the CA1 region, were randomly imaged. Confocal *z*-stacks covering the whole depth of the neuron somata were taken with 0.38 μm step size using a 63× oil objective with 2× digital zoom magnification (Leica TCS SPE, Germany). Stacks were then processed using FIJI/ImageJ software. Single confocal planes from each neuron, in which the penetration of each antibody was optimal, were selected. Briefly, the profile of the soma of these neurons was drawn manually and the selection was enlarged 1.25 μm to cover the area surrounding the somata. The region of interest (ROI) was then binarized and puncta was defined as a structure displaying an area not smaller than 0.15 μm^2^ and not larger than 2.5 μm^2105^. The density of puncta (number of puncta *per* micron of soma perimeter) was analyzed as described above (see “Analysis of the excitatory/inhibitory balance”). The percentage of area covered by PV+ puncta was additionally analyzed in P90 mice.

### Analysis of PolySia-NCAM Immunoreactivity

Sections starting at Bregma −1.94 mm were selected to study the immunoreactivity of polySia-NCAM in the different layers of the hippocampal CA1 region as described previously^106^. Samples were examined with an Olympus CX41 microscope under bright-field illumination, homogeneously lighted and digitized using a CCD camera. Photographs of the different layers were taken randomly at 20x magnification using the same exposure time and ISO. Grey levels were converted to optical densities using FIJI/ImageJ software (NIH) and normalized with respective internal white matter regions.

### Quantification of PVALB Cells, PNNs and Analysis of Hippocampal Volume

To study the number of PVALB neurons (PV+), PNNs and their co-localization in CA1 hippocampal region, we used a modified version of the fractionator method^107,108^. Briefly, within each 50-µm-thick section of one from the six systematic-random series of sections, all labeled cells covering the 100% of the sample area were counted. Confocal *z*-stacks covering the whole depth were taken with a confocal microscope (Olympus FV-10) using a 10× objective to obtain the 2D projections. The images were processed afterward using FIJI/ ImageJ software^99^.

A volumetric analysis of the hippocampal CA1 region was also performed on these sections. Microphotographs at 10x were obtained with a confocal microscope, and then processed using the stitching plugin in FIJI/ImageJ^99^ in order to have the whole nucleus on a single microphotograph. All the sections in a 1/6 subseries containing the region of interest were analyzed and the areas estimated using Cavalieri’s principle^109^. To obtain the volume of the CA1 region, we multiplied the area by the thickness of the slice (50-µm) and the number of series. We finally summed up the volumes across all slices for each animal.

### Expression of EPOR in GABAergic Neurons

RNAscope Multiplex Fluorescent assay v2 (CatNo. 323100) from Advanced Cell Diagnostics (ACD; Hayward, CA, USA) was used for the detection of *Epor* mRNA in GAD67 and PVALB positive cells, encoded by *Gad1* and *Pvalb* genes, respectively. C57BL/6 mice were perfused intracardially using 0.9% NaCl, and their brains were post-fixed using 4% paraformaldehyde for 24 hours. The brains were then sectioned into 50 µm-thick slides using a freezing sliding microtome (Leica SM2010 R) and two hippocampi *per* mouse were mounted on SuperFrost Plus slides (CatNo. 12-550-15, Fisherbrand). After that, the RNAscope assay was performed following manufacturer’s instructions. In short, the slides were dehydrated, exposed to hydrogen peroxide solution (10min., CatNo. 322335, ACD) and target retrieval solution (15 min., CatNo. 322000, ACD) at 98°C-102°C. Then, the slides were incubated in RNAscope Protease III (30 min., CatNo. 322337, ACD) at 40°C using the HybEZ II hybridization system (CatNo. PN 321710, ACD), a temperature that was kept constant for all incubation steps during the rest of the assay. The samples were then incubated for 2 hours using the corresponding probes to detect *Epor* (Mm-Epor, CatNo. 412351, ACD), *Gad1* (Mm-Gad1, CatNo. 400951-C2, ACD) as a general interneuronal marker, and *Pvalb* (Mm-Pvalb, CatNo. 421931-C3, ACD) as one of the main interneuronal subpopulation. Afterwards, the probes were amplified using AMP1 (30 min., CatNo. 323101, ACD), AMP2 (30 min., CatNo. 323102, ACD), and AMP3 (15 min., CatNo. 323103, ACD), followed by the development of each probe. First, we developed the *Epor* signal incubating the samples with HRP-C1 (30 min., CatNo. 323104, ACD) and Opal 570 (30 min., CatNo. FP1487001KT, Akoya biosciences), followed by the blockade of the horseradish peroxidase action using HRP-blocker (CatNo. 323107, ACD). We then developed the *Gad1* signal, this time using HRP-C2 (30 min., CatNo. 323105, ACD), Opal 520 (30 min., CatNo. FP1487001KT, Akoya biosciences) and HRP-blocker. Finally, we developed *Pvalb* signal using HRP-C3 (30 min., CatNo. 323106, ACD), Opal 690 (CatNo. FP1487001KT, Akoya biosciences) and HRP-blocker. Samples were coverslipped using fluorescence mounting media (CatNo. S302380-2, Agilent). Additionally, positive and negative controls were also added to the assay and followed the same protocol as the rest of the samples (3-plex positive control probe Mm, CatNo. 320881, and negative control probe, CatNo. 320871 from ACD). To estimate the percentages of interneurons that expressed EPOR, all hippocampi were imaged (2 *per* animal), focusing on the dorsal CA1 region. We used a laser scanning confocal microscope (Leica TCS SPE) with a 63x immersion objective and a 2.5x additional digital zoom to cover the whole area. The percentage of *Epor* expression was finally calculated in the general population of interneurons (*Gad1+*), or specifically in parvalbumin (*Pvalb+*) cells, always considering the animal as the sampling unit.

### NanoSIMS

After fluorescence immunohistochemistry of the sections, CA1 hippocampal regions were dissected and embedded in LR-White-Resin (AGR1281; Agar Scientific, Wetzlar, Germany) and accelerator mixture (AGR1283; Agar Scientific) following a dehydration protocol as described previously^70^. Briefly, 30, 50 and 70% ethanol was applied to each section in successive 10-min steps. After drying, sections were brought into a solution containing 70% ethanol and LR-White-Resin. To proceed with resin polymerization, sections were surrounded by a silicon cylinder, and pure LR-White-Resin with accelerator mixture were brought inside of the cylinder. Polymerization was finished after 90 min incubation at 60°C. Sections were further cut into 200 nm thick sections with an EM-UC6 Ultramicrotome (Leica Microsystems) and placed on silicon wafers (Siegert Wafer GmbH, Aachen, Germany). Epifluorescent imaging of PV+ expression with a Nikon Ti-E inverted microscope and 100x objective (NA 1.59) was followed by scanning with a Cs+ primary ion beam in a nanoscale secondary ion mass spectrometry (NanoSIMS 50L) instrument (Cameca, Gennevilliers Cedex, France). Samples were eroded and ionized at 60 pA for 3min, and the measurements were carried out at 2.5 pA with a dwell time per pixel of 4000 ms. From the resulting secondary ions, ^12^C^14^N- and ^12^C^15^N-were detected and measured, and are referred to as ^14^N and ^15^N, respectively, in this work. The mass resolving power was tuned to enable optimal separation of ^12^C^15^N-from ^13^C^14^N. For each measurement, 3 planes of 40×40 mm (256×256 pixels) were recorded in a total of 4 animals/group, drift-corrected, and summed for analysis using OpenMIMS-plugin (NRIMS) for ImageJ. The resulting NanoSIMS images were then aligned to corresponding fluorescent images in Adobe-Photoshop. PV-positive signals were selected and corresponding NanoSIMS regions quantified in MATLAB (Mathworks, Ismaning, Germany) using a custom written plugin. PV-positive regions were normalized to negative ones and compared between conditions in GraphPad Prism9.

### Single Nuclei RNA-Sequencing

On P49, 24h after the last PL/rhEPO injection, 23 mice (n=11 rhEPO, n=12 PL) were sacrificed by cervical dislocation. The brain was immediately removed, the right hippocampus dissected on an ice-cold plate, quickly immersed in liquid nitrogen and kept at −80°C. Two right hippocampi of the same treatment group were collected in one tube for sequencing (one tube in the rhEPO group with only one right hippocampus). Final analysis was performed on n=6 tubes *per* group. Single nucleus suspension was prepared using 10x Genomics Chromium Single Cell 3’ Reagent Kits v3 (10X Genomics, Pleasanton, CA) according to manufacturer’s protocol. Quantity and quality of cDNA were assessed by Agilent 2100 expert High Sensitivity DNA Assay. cDNA samples were sequenced on NovaSeq 6000 S2 flowcell at UCLA Technology Center for Genomics and Bioinformatics. Raw and processed snRNA-seq data are publicly available on GEO via accession code GSE220522.

We commenced our data analysis by reanalyzing the recently published datasets from our laboratory^21^. Briefly, the raw sequences were aligned to the mouse genome (mm10) using 10x Genomics CellRanger software. The alignment was run with standard parameters described in the developer’s manual. Afterwards, any potential issues arising from the sequencing technology were resolved using CellBender and CellRanger software^110^. We then employed Seurat (v4.1.1)^111^ implemented in R (v4.1.0) for filtering, normalization, and cell-types clustering. Data were normalized by regressing the impact of mitochondrial, cell-cycle, and ribosomal genes. Cell types were clustered using the first 30 PCA dimensions that were further fed in constructing the shared-nearest neighbor (SNN) graph. Major cell types were assigned based on the popular markers, and cell subtypes within major cell types were annotated using the sub-cluster markers obtained using default parameters.

To dissect the interneuronal lineages, we chose the cell-type cluster labelled Interneurons^21^, which was validated by testing the co-expression of the *Gad1* and *Gad2* genes. Additionally, we also tested whether the transcriptional markers of the rest of the hippocampal lineages were absent to determine their fidelity as interneurons. To classify these diverse lineages of interneurons, we utilized the set of known bonafide markers. From this data, we used only picked predefined lineages of interneurons for further analysis to perform a comparative analysis between rhEPO and PL samples.

Furthermore, we used the WebGestalt tool^112^ to find the over-representation of gene ontologies within the differentially expressed genes between rhEPO and PL lineages. We finally used the ‘Liana’ tool^55^ to decipher the ligand-receptor dynamics within the Interneuron lineages and compared them between rhEPO and PL samples. All the graphics displayed in this manuscript are built on the R platform. Specific codes of data/plots and the way lineages and cell types are classified in our study are available on GitHub link (https://github.com/Manu-1512/Yasmina-et-al.-Interneurons).

### Statistical Analysis

Data is shown as mean ± standard error of the mean (SEM), with N numbers (i.e., number of mice/group) and statistical test specified in the text or corresponding figure legend. All statistical analyses and graphs were performed with GraphPad Prism 9 software. Normality and homoscedasticity of data were initially evaluated with Kolmogorov-Smirnov and Levene’s tests, respectively. Student’s unpaired two-tailed t tests were performed in normally distributed data and two-tailed Mann-Whitney U test in non-parametric data. Probability values lower than 0.05 (*p* < 0.05) were considered as statically significant.

## ACKNOWLEDGEMENTS

This work has been supported by the European Research Council (ERC) Advanced Grant to HE under the European Union’s Horizon Europe research and innovation programme (acronym *BREPOCI;* grant agreement No 101054369), as well as by the Max Planck Society, the Max Planck Förderstiftung, the Deutsche Forschungsgemeinschaft (DFG, German Research Foundation), via DFG-Center for Nanoscale Microscopy & Molecular Physiology of the Brain (CNMPB). Research in the labs of HE and KAN is supported by DFG TRR-274 /1 2020 – 408885537. YC is recipient of a grant from the Peter and Traudl Engelhorn Foundation. HC is supported by a postdoctoral “Margarita Salas” (MS21-074) grant from the Universitat de València funded by the Spanish Ministry of Science and the Next Generation EU. Research at VT lab is supported by a project (PID2019-108562GB-I00) funded by MCIN/AEI/ 10.13039/501100011033 and by “ERDF A way of making Europe”. Research in the labs of DG, RK and KAN is supported by the Adelson Medical Research Foundation. Research in JN lab is supported by a project (PID2021-127595OB-I00) funded by MCIN/AEI/ 10.13039/501100011033 and by “ERDF A way of making Europe” and the Generalitat Valenciana (PROMETEU/2020/024).

## AUTHOR CONTRIBUTIONS

Concept, design, supervision: YC, JN, HE, MS, VT

Drafting manuscript and display items: YC, MS, JN, HE

Data acquisition/generation: YC, HC, PK, MPR, QW, KG, RK, MS

Data analyses & interpretation: YC, MS, VT, RK, DG, KAN, SR, JN, HE

**All authors read and approved the final version of the manuscript.**

---

**Supplementary Figure 1.**
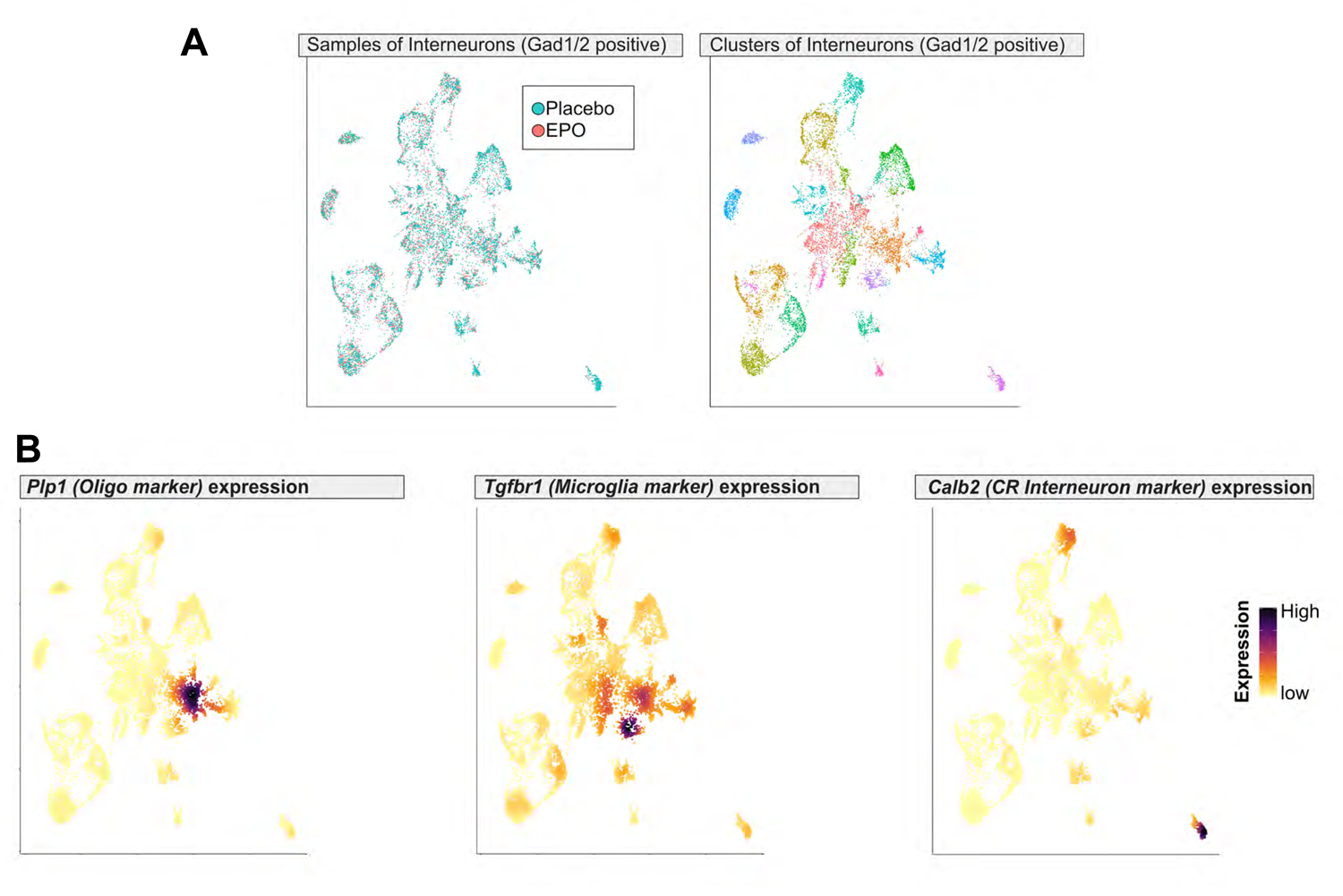
**A:** UMAP biplot illustrates the re-clustering of interneuron lineages (high *Gad1* and *Gad2* expression) filtered from the whole hippocampus snRNA-seq data^21^. Left panel shows that rhEPO and PL samples contributed similarly in each cluster. Right panel demonstrates the different clusters within interneuron lineages obtained using default features of Seurat algorithm. Every dot is a single nucleus, and colors denote the distinct clusters. **B:** Multiple feature plots based on UMAP of interneuron clusters in rhEPO and PL samples displaying unsupervised identification of expression markers of oligodendrocytes (*Plp1*), microglia (*Tgfbr1*), and Cajal-Retzius cells (*Calb2*). Dots in gold/maroon denote lower/higher expression in each single nucleus, respectively.

**Supplementary Figure 2.**
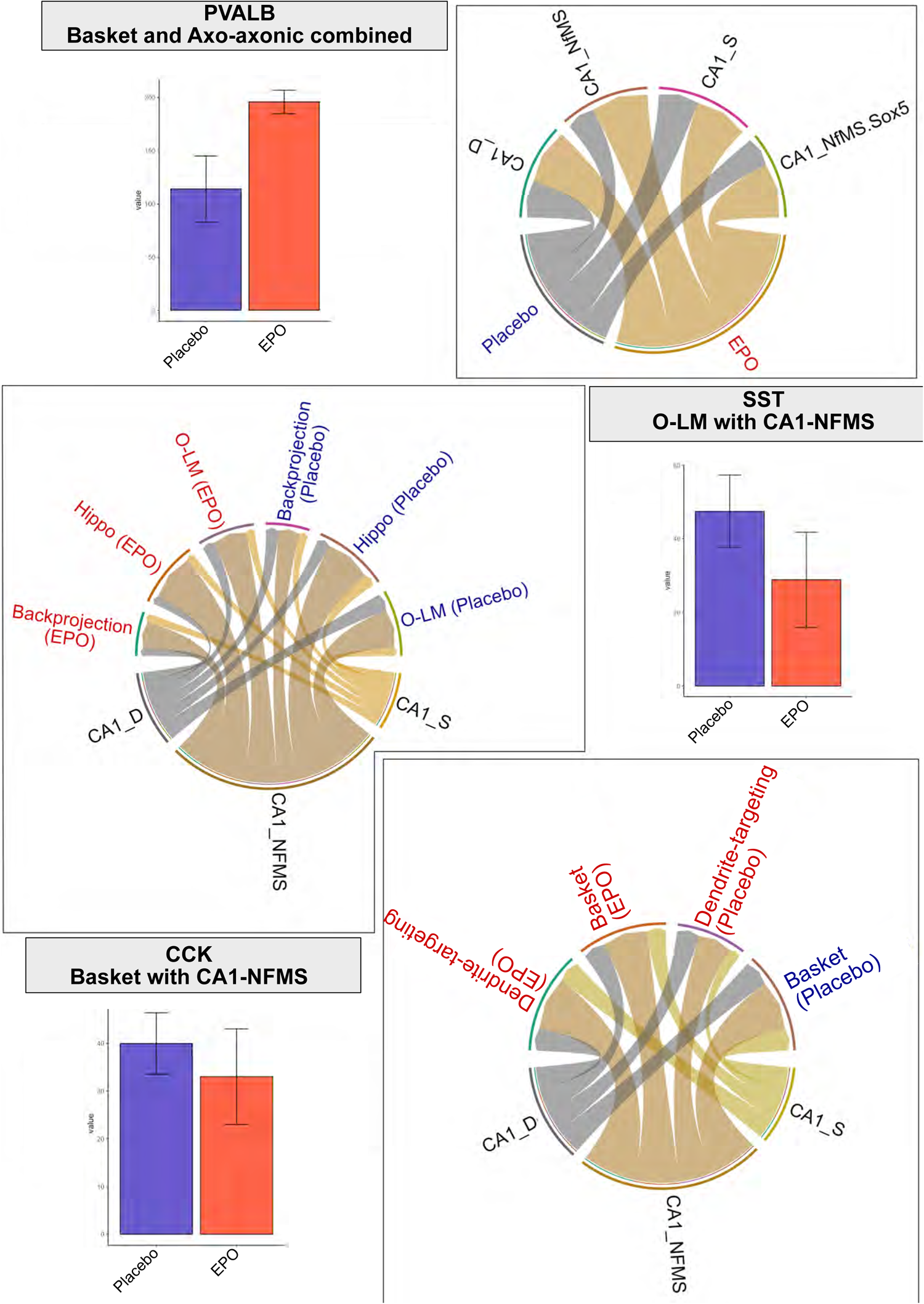
The global interaction between interneuron and pyramidal lineages is both data-driven and database-curing and illustrates the differential interaction between rhEPO and PL samples. Receptor/ligand or sender/receiver cross-talk pattern between cell types of interneurons and pyramidal layers are visualized in a circular plot where the source cell type (ligands) is connecting with the targets (receptors). To show an example of the empirical estimation of the number of interactions, a few selected barplots are displayed next to circos plots. Blue corresponds to PL, red to rhEPO. Cell types are explained in Singh et al 2023^21^: CA1_NFMS= Newly formed-migrating-superficial, CA1_S= superficial, CA1_D= dorsal and CA1_NfMS.Sox5= Newly formed-migrating-superficial-Sox5 pyramidal neurons from the CA1 region.

