## Supplemental table 4 for "Erythropoietin restrains the inhibitory potential of interneurons in the mouse hippocampus"

| Anti | Host | Isotype/Label | Dilution | Incubation | Company |
|---|---|---|---|---|---|
| **PRIMARY ANTIBODIES** | | | | | |
| GFP | Chicken | Ig Y | 1:1000 | 48h, 4°C | Abcam |
| PV | Guinea Pig | Ig G | 1:2000 | 48h, 4°C | Synaptic Systems |
| CaMKII-α | Mouse | Ig G1 | 1:500 | 48h, 4°C | Abcam |
| CB1r | Rabbit | Ig G | 1:1000 | 48h, 4°C | Synaptic Systems |
| Wisteria Floribunda Lectin Biotin | | | 1:200 | 48h, 4°C | Sigma-Aldrich |
| VGLUT1 | Guinea Pig | Ig G | 1:2000 | 48h, 4°C | Millipore |
| VGAT | Rabbit | Ig G | 1:1000 | 48h, 4°C | Synaptic Systems |
| PSA-NCAM | Mouse | Ig M | 1:1400 | 48h, 4°C | DSHB |
| GAD67 | Mouse | Ig G2a | 1:500 | 48h, 4°C | Sigma-Aldrich |
| PV | Mouse | Ig G1 | 1:500 | 48h, 4°C | Sigma-Aldrich |
| **SECONDARY ANTIBODIES** | | | | | |
| Chicken IgY | Goat | Alexa 488 | 1:400 | 2h, 25°C | Life Technologies |
| Guinea pig IgG | Goat | Alexa 555 | 1:400 | 2h, 25°C | Life Technologies |
| Mouse IgG1 | Goat | Alexa 555 | 1:400 | 2h, 25°C | Life Technologies |
| Guinea Pig IgG | Goat | DyLight 649 | 1:400 | 2h, 25°C | Jackson ImmunoResearch |
| Rabbit IgG | Donkey | Alexa 647 | 1:400 | 2h, 25°C | Molecular Probes |
| Avidin | | Alexa 635 | 1:400 | 2h, 25°C | Invitrogen |
| Rabbit IgG | Goat | Alexa 635 | 1:400 | 2h, 25°C | Life Technologies |
| Guinea Pig IgG | Donkey | Cy3 | 1:500 | 2h, 25°C | Jackson ImmunoResearch |
| Mouse IgM | Donkey | Biotin | 1:400 | 2h, 25°C | Jackson ImmunoResearch |
| Mouse IgG | Horse | Biotin | 1:200 | 1.5h, 25°C | Vector Laboratories |

**Table S4.** Primary and secondary antibodies used in the study.
